# Extensive and accurate benchmarking of DIA acquisition methods and software tools using a complex proteomic standard

**DOI:** 10.1101/2020.11.03.365585

**Authors:** Clarisse Gotti, Florence Roux-Dalvai, Charles Joly-Beauparlant, Loïc Mangnier, Mickaël Leclercq, Arnaud Droit

**Affiliations:** Proteomics platform, CHU de Québec - Université Laval Research Centre, Québec City, Québec, Canada; Computational Biology Laboratory, CHU de Québec - Université Laval Research Centre, Québec City, Québec, Canada

**Keywords:** Proteomic standard, software benchmarking, data-independent acquisition, label-free quantification

## Abstract

Over the past decade, the data-independent acquisition mode has gained popularity for broad coverage of complex proteomes by LC-MS/MS and quantification of low-abundance proteins. However, there is no consensus in the literature on the best data acquisition parameters and processing tools to use for this specific application. Here, we present the most comprehensive comparison of DIA workflows on Orbitrap instruments published so far in the field of proteomics. Using a standard human 48 proteins mixture (UPS1 – Sigma) at 8 different concentrations in an *E. coli* proteome background, we tested 36 workflows including 4 different DIA window acquisition schemes and 6 different software tools (DIA-NN, DIA-Umpire, OpenSWATH, ScaffoldDIA, Skyline and Spectronaut) with or without the use of a DDA spectral library. Based on the number of proteins identified, quantification linearity and reproducibility, as well as sensitivity and specificity in 28 pairwise comparisons of different UPS1 concentrations, we summarize the major considerations and propose guidelines for choosing the DIA workflow best suited for LC-MS/MS proteomic analyses. Our 96 DIA raw files and software outputs have been deposited on ProteomeXchange for testing or developing new DIA processing tools.

## INTRODUCTION

Over the past two decades, proteomics, or analysis of the protein content of a biological sample, has become an essential strategy for the comprehension of systems biology and molecular events involved in health and disease. *Bottom-up proteomics*, which relies on liquid chromatography-tandem mass spectrometry (LC-MS/MS) to analyze peptides resulting from proteolysis, allows the identification and quantification of up to thousands of proteins in a whole cell extract^1,2^.

The most widely used strategy to obtain broad coverage of the proteome, known as *discovery proteomics*, is based on data-dependent acquisition (DDA), in which MS1 survey scans containing accurate mass measurement of co-eluting peptides are acquired successively, followed by acquisition of fragmentation (or MS2) spectra of automatically selected precursor ions^3^. MS1 and MS2 spectra are used to query publicly available protein databases in order to obtain peptide identification and then infer the corresponding proteins. Moreover, quantification information can be obtained using, for instance, the area under peaks reconstructed from MS1 precursor ion intensities by *label-free quantification* (LFQ)^4^. The DDA method allows thousands of proteins to be identified and quantified relatively in fractionated or purified or whole-protein extracts. This large-scale analysis provides essential information on biological mechanisms.

In *targeted proteomics*, a previously chosen list of proteins of interest is analyzed in a protein extract using selected reaction monitoring (SRM)^5,6^ or parallel reaction monitoring (PRM)^7^, by successive measurements of specific precursor/fragment transitions of a given peptide throughout the chromatographic elution. In this case, chromatographic elution profile overlap of several transitions of a same peptide validates detection, and the peptide is quantified by integrating the area under the chromatographic peak. Compared to *discovery proteomics*, sensitivity, quantification accuracy and reproducibility are improved across numerous samples. For these reasons, the targeted approach is well suited to monitoring protein candidates in large sample cohorts in biomarker validation studies.

Although these two approaches have been adopted widely, both have drawbacks. Protein identification by DDA is based on a stochastic precursor selection event prior to the acquisition of fragmentation spectra, leading to a lack of run-to-run reproducibility when complex samples are analyzed. Although post-acquisition bioinformatics tools are now better able to deal with this under-sampling effect by recovering the precursor MS1 signal in all runs of an experiment, the dynamic range of intensities in MS1 spectra is broad and often limits detection and quantification of low-abundance species. In contrast, by narrow quadrupole filtering of the targeted precursor, SRM or PRM analyses generally avoid the concomitant selection of abundant species, making targeted proteomics more sensitive and more reproducible than DDA. However, even when SRM/PRM transitions signal acquisition is finely scheduled over the whole chromatographic run time, this strategy is limited to the analysis of few hundreds of peptides due to the number of transitions to be monitored in parallel without exceeding the appropriate cycle time for chromatographic peak reconstruction.

To overcome these limitations, an alternative *discovery proteomics* strategy has been devised, combining the strengths of the DDA and targeted approaches to obtain reproducible and accurate quantification at proteome level (several thousands of proteins). Developed at the beginning of the millennium^8–10^ and popularized by Gillet *et al.^11,12^*, *data-independent acquisition* or DIA relies not on precursors selected individually, but on systematic windows of precursors and fragmentation of all peptide ions contained in each window. The main advantages of this strategy are the coverage of all detectable ions present in a sample and a smaller dynamic range of intensities within the spectra used for quantification. Indeed, DIA quantification uses MS2 spectra resulting from small precursor windows rather than acquisition over the full mass range as in MS1 spectra used for quantification in DDA. This makes the quantitative aspect of DIA analysis more reproducible than DDA, especially for low-abundance species, while identifying thousands of proteins^12,13^. However, the resulting fragmentation spectra made highly complex by selection of multiple precursors makes database search engines usually used for DDA analysis, unable to match experimental data to theoretical masses obtained from the public protein databases. To identify peptides in chimeric DIA-MS2 spectra, two main strategies have been devised: spectrum-centric and peptide-centric. In spectrum-centric strategy, identifications are obtained by the deconvolution of the complex spectra and search against protein databases, whereas in peptide-centric identification, m/z and retention time information, in previously generated spectral libraries, are used to extract signal from DIA spectra^14^. In recent years, several open-source or proprietary software tools have been developed for DIA analysis^15–20^. However, each has its own specificity, able to process data using a DDA spectral library and/or a FASTA file only, with or without normalization and missing values imputation algorithms, with or without a graphical user interface^21^. The DIA window acquisition scheme appears to be a critical point for DIA analysis since wide windows allow coverage of the whole mass range in a reasonable duty cycle time whereas narrow windows produce simpler spectra, facilitating subsequent analysis. Moreover, overlapping windows have been shown to increase the number of protein identifications while maintaining the proper cycle time for the quantification of each analyte^22^ and several studies mentioned the use of variable window sizes^19,23,24^. Among the plethora of methods for DIA acquisition and processing mentioned in the literature, the user may find it difficult to choose the best strategy for complex proteomes analysis and especially for the detection and accurate quantification of less abundant species. In 2016, in a collaboration of software developers, several DIA tools were compared using a proteomic dataset acquired from a mixture of 3 protein extracts from human cells, yeast, and *Escherichia coli* in SWATH-MS DIA mode on TripleTOF (Sciex) instruments^23^. Although the experimental design allowed relative quantification through measurement of different protein ratios for each organism, the concentration of each protein in the mixture remained unknown. It was not possible to draw any conclusions about software performance in relation to protein abundance in the sample. Since this study was published, new tools have been developed and the capabilities of others have been extended to library-free searches. In the present study, we compared the quantitative performance of DIA-NN, DIA-Umpire, OpenSWATH, ScaffoldDIA, Skyline and Spectronaut using a proteomic standard composed of 48 human proteins (a commercial mixture called UPS1, Sigma-Aldrich) spiked in a whole *E. coli* protein extract background. A similar proteomic standard has been used previously to test DDA spectral processing tools^25–27^. Data obtained at 8 UPS1 concentrations on an Orbitrap Fusion (Thermo) instrument with 4 different DIA window schemes (narrow, wide, mixed, overlapped) were analyzed and processed with or without a large spectral library containing about 2800 *E. coli* and UPS1 proteins. We thus compiled number of identified and quantified proteins, quantification linearity and reproducibility, ratio accuracy, sensitivity, and specificity in pairwise comparisons using a total of 36 acquisition and processing combinations: the 4 DIA window schemes with 4 DIA tools using a spectral library and with 5 DIA tools using a FASTA file (Fig. 1). Our goal was to identify the best quantification strategy for proteins at different abundances in complex proteomes. Our 96 DIA raw data files, spectral library, intermediate files, software outputs and quantification tables may be accessed via ProteomeXchange for further statistical analyses or software testing and development.

**Figure 1.**
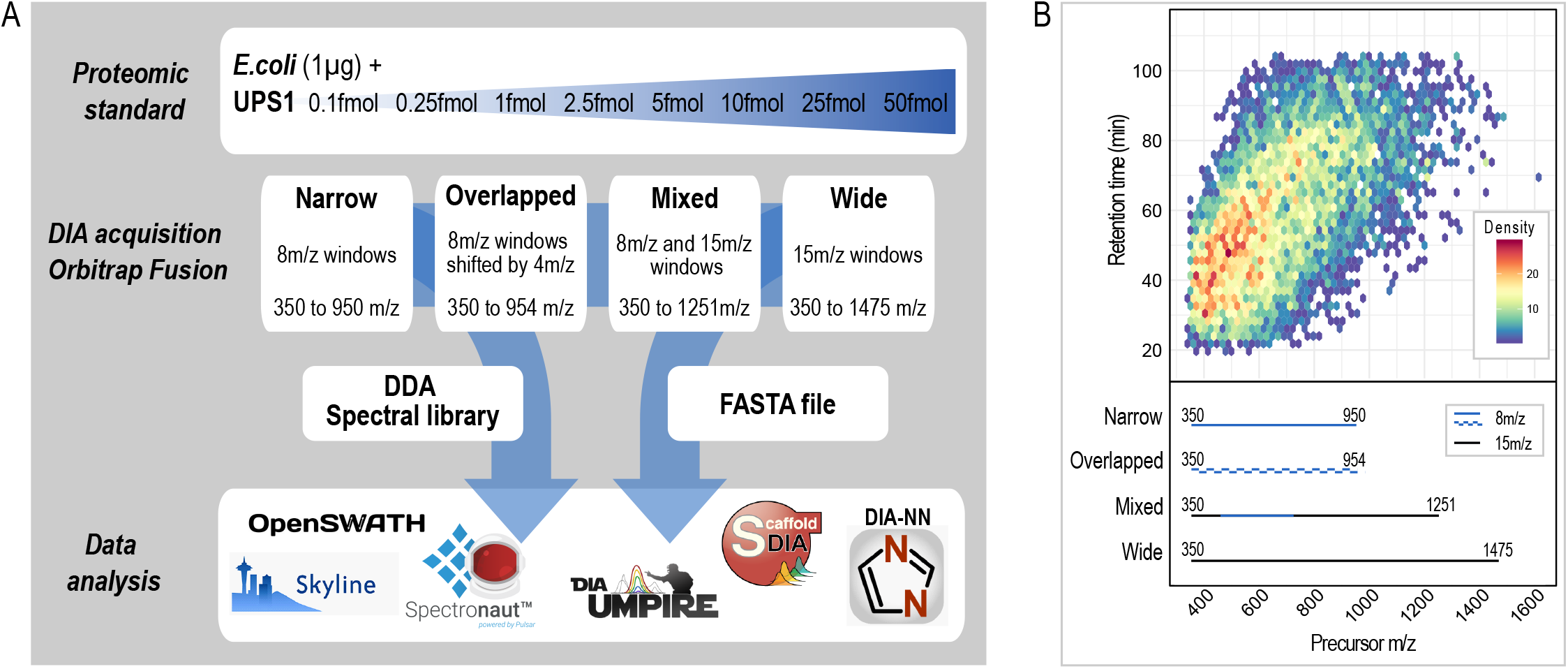
Proteomic standard and processing workflows. (A) The proteomic standard, comprising UPS1 (Sigma) at 8 concentrations in *E.coli* protein extract background, was analyzed by LC-MS/MS with 4 different DIA acquisition schemes. Raw data files were processed using 1 of 6 different software tools and a FASTA file only or/and a DDA spectral library. (B) Distribution of precursor ions from a sample containing 200 fmol of UPS1 per microgram of *E.coli*, over the mass range and run time (DDA mode). Window schemes for DIA analysis are shown below the graph.

## METHODS

### Sample preparation

#### Bacterial culture and protein extraction

*Escherichia coli* (#CCRI-12923) was obtained from the Collection du Centre de Recherche en Infectiologie, Université Laval (CCRI, Québec, Canada). It is registered as WDCM861 at the World Data Centre for Microorganisms. Six aliquots (1 mL) broth culture in Brain Heart Infusion (BHI) medium at a concentration of (8×10^8^ cfu/mL) were centrifuged at 10,000 × *g* for 15 min and the pellets were washed three times with 1 mL of 50 mM Tris. Dried pellets were frozen and stored at −20°C.

For protein extraction, each aliquot was resuspended in 135 μL of extraction buffer (50 mM ammonium bicarbonate, 1% sodium deoxycholate and 20 mM 1,4 dithiothreitol. Bacteria were then inactivated by heating for 10 min at 95°C and lysis was achieved by sonication with Bioruptor® (Diagenode) for 15 minutes with 30s/30s ON/OFF cycles at high intensity.

Lysed cells were centrifuged at 13,000 × *g* for 10 min to remove debris, the 6 supernatants were then pooled, and the protein concentration was measured using the Bradford assay. Protein extracts were stored at −80°C.

#### Proteomic standard

Universal Proteomic Standard-1 (UPS1, Sigma) containing 48 human proteins (5 pmol each) was diluted serially in *E. coli* whole protein extract (previously diluted at 0.1 μg/μL in extraction buffer) to obtain 8 different UPS1 concentrations: 50, 25, 10, 5, 2.5, 1, 0.25 and 0.1 fmol of UPS1 per μg of *E. coli.* Disulfide bridges were reduced by heating for 30 minutes at 37°C and alkylated with 50 mM iodoacetamide for 30 minutes in darkness. Prior to trypsin (Promega) digestion, the pH was adjusted to 8.0 with sodium hydroxide. Proteolysis was carried out overnight at an enzyme/protein ratio of 1:50 at 37°C. The reaction was stopped by dropping the sample pH to 2.0 with formic acid. After centrifugation at 16,000 × *g* for 5 minutes to remove the deoxycholate precipitate, the supernatants (peptides) were cleaned on an Oasis HLB cartridge 10 mg (Waters) according to the manufacturer instructions, aliquoted and dried under vacuum.

#### Spectral library

To generate a spectral library, 250 μg of *E. coli* protein extract were used. Reduction, alkylation, and trypsin digestion were performed under the conditions described above. Peptides were fractionated on an Agilent 1200 Series HPLC system equipped with an Agilent Extend C_18_ (1.0 mm × 150 mm, 3.5 μm) column. Peptides were loaded at 1 mL/min of solvent A (10 mM ammonium bicarbonate, pH 10) and eluted by addition of solvent B (90% acetonitrile/10% ammonium bicarbonate pH 10) with a gradient of 5–35% B for 60 minutes and 35–70% B for 24 minutes. Fractions were collected in a 96-well micro-assay plate every 1 minute and finally pooled into 48 fractions and vacuum-dried.

A sample of 200 fmol UPS1 per μg of *E. coli* extract was prepared under the same conditions to complete the spectral library with UPS1 human proteins.

### Data acquisition

#### NanoLC-MS/MS analysis

Samples were resuspended at 0.2 μg/μL in loading buffer (2% acetonitrile/0.05% trifluoroacetic acid) containing 0.5X iRT peptides (Biognosys). Injected volume was 5 μL. The nano-LC-MS/MS system comprised a U3000 NanoRSLC liquid chromatography system (ThermoScientific, Dionex Softron GmbH, Germering, Germany) in line with an Orbitrap Fusion Tribrid – ETD mass spectrometer (ThermoScientific, San Jose, CA, USA) driven by Orbitrap Fusion Tune Application 3.0.2041 and equipped with a Nanospray Flex™ ion source. Peptides were trapped at 20 μL/min in loading solvent (2% acetonitrile/0.05% trifluoroacetic acid) on a 300 mm i.d × 5 mm, C_18_ PepMap100, 5 mm, 100 Å precolumn cartridge (Thermo Fisher Scientific) for 5 minutes. The pre-column was switched in line with a PepMap100 RSLC, C_18_ 3 mm, 100 Å, 75 μm i.d. × 50 cm length column (Thermo Fisher Scientific) and the peptides were eluted with a linear gradient from 5–40 % solvent B (A: 0.1% formic acid, B: 80% acetonitrile/0.1% formic acid) in 90 minutes, at 300 nL/min. Spectra were acquired using Thermo XCalibur software version 4.1.50. Lock mass internal calibration on the m/z 445.12003 siloxane ion was used.

#### DDA acquisition parameters – Spectral library

Full-scan mass spectra (350–1800 m/z) were acquired in the Orbitrap using an AGC target of 4e5, a maximum injection time of 50 ms and a mass resolution of 120,000. Each MS scan was followed by acquisition of fragmentation MS/MS spectra of the most intense ions for a total cycle time of 3 seconds (top speed mode). The selected ions were isolated using the quadrupole analyzer in a window of 1.6 m/z and fragmented by higher energy collision-induced dissociation (HCD) with 35% collision energy. The resulting fragments were detected in the Orbitrap at a 15,000 resolution with an AGC target of 5e4 and a maximal injection time of 22 ms. Dynamic exclusion of previously fragmented peptides was set for a period of 30 sec and a tolerance of 10 ppm.

#### DIA acquisition parameters – Proteomic standard

Full-scan mass spectra were acquired in the Orbitrap using an AGC target of 4e5, a maximal injection time of 50 ms and a resolution of 60,000. Each MS scan was followed by 75 DIA scans with different isolation window widths depending on each scheme: 8 m/z windows spanning the 350–950 precursor mass range for *Narrow*, two cycles of 8 m/z isolation windows shifted by 4 m/z spanning the 350–954 range for *Overlapped*, a combination of 8 m/z for 455–711 and 15 m/z for 350–455 and 711–1251 for *Mixed,* and 15 m/z windows spanning 350–1475 for *Wide* (Fig. 1 and Supplementary Table S1). Precursor ions were selected in the quadrupole, fragmented by HCD with 35% collision energy and fragments were detected in the Orbitrap at a resolution of 15,000 and with an AGC target of 4e5 and a maximal injection time of 22 ms.

### Data processing and statistics

#### Spectral library generation

Raw data from *E. coli* fractionation and 200 fmol/μg UPS1 DDA acquisition were searched against an *E. coli* database (UniProt Reference Proteome – Taxonomy 83333 – Proteome ID UP000000625 – 4312 entries – 2016.03.15) and to the UPS1 database (downloaded from Sigma – 48 entries), using Mascot search engine version 2.5.1 (Matrix Science) as a node in Proteome Discoverer 2.3.0.523. The trypsin enzyme parameter was set for two possible missed cleavages. Carbamidomethylation of cysteines was set as a fixed modification and methionine oxidation was set as a variable modification. Mass search tolerances were 10 ppm and 25 mmu for MS and MS/MS respectively. Peptide and protein identifications were filtered at 1% False Discovery Rate (FDR).

Mascot *.dat* files were then used to generate a spectral library in Skyline software^28^ (version 20.2.0.343) through a *.blib* file. Another spectral library was generated in Spectronaut v14.10.20122 (Biognosys AG)^19^ using the *.pdresults* file of Proteome Discoverer. Detailed parameters used to generate both spectral libraries are listed in Table S1.

#### DIA file processing

The tools DIA-NN^18^ v02/04/2020, DIA-Umpire v2.0^17^, OpenSWATH through diaproteomics workflow v.1.1.0^15,29^, ScaffoldDIA v2.1.0 (Proteome Software, Inc.)^20^, Skyline v20.2.0.343^28^, and Spectronaut v14.10.20122 (Biognosys AG)^19^ were used for extracting peptide signals from raw files using the previously generated spectral library (DIA-NN, OpenSWATH, ScaffoldDIA, Skyline, Spectronaut) (*Library* mode) or using a single *E. coli* and UPS1 FASTA file (UniProt Reference Proteome – Taxonomy 83333 – Proteome ID UP000000625 – 4312 entries – 2016.03.15 and the 48 sequences of UPS1) (DIA-NN, DIA-Umpire, ScaffoldDIA, Spectronaut) (*FASTA* mode). The tools in the two processing modes (*Library* or *FASTA*) were used as recommended by the user manual or by the software developers (Table S1).

Raw files were converted to *.mzML* files for analysis by DIA-NN, OpenSWATH and Skyline or to *.mzXML* files for analysis by DIA-Umpire. ScaffoldDIA and Spectronaut use the raw files as input and convert them with their own embedded tool (Table S1).

For the *Library* mode, Spectronaut and Skyline were used with their own DDA spectral libraries, while the Skyline *.blib* file was also used in ScaffoldDIA. The precursor/fragment transitions corresponding to this *.blib* file were also exported in Skyline as a *.tsv* file to be used in DIA-NN and OpenSWATH.

For analysis in *FASTA* mode, a DIA spectral library was generated from the DIA files in DIA-NN or DIA-Umpire prior to signal extraction and quantification. For ScaffoldDIA, an *in-silico* Prosit library was generated and used for the processing of DIA files. Spectronaut was used in ‘DirectDIA’ mode. (Table S1).

The following settings were common to all software: maximum of 2 missed trypsin cleavages, cysteine carbamidomethylation as fixed modification and methionine oxidation as variable modification, a minimum of 4 fragments and a maximum of 6 were used to consider a peptide for quantification and only peptides with charges of 2+ to 4+ were saved. Data were filtered out at a false discovery rate (FDR) of 1% or at *q* value < 0.01 at precursor, peptide and/or protein level depending on the capabilities of each tool. The specific parameter settings for each software tool are detailed in Table S1.

#### Data post-processing and evaluation of DIA label-free quantification

All data post-processing and visualization was performed using R software^30^ (Fig. S1). After removing contaminants and decoy features, precursor table output of each tool corresponding to the validated proteins and peptides were used for subsequent analysis. OpenSWATH and Spectronaut reported a few extremely low outliers that were obviously extraction errors and were removed using an arbitrary intensity cut-off of 10 for Spectronaut and 1 for OpenSWATH. Since DIA-NN, ScaffoldDIA and Spectronaut perform their own data normalization, the normalized precursor intensity values were used. For the other tools (DIA-Umpire, OpenSWATH and Skyline) precursor intensity values were normalized by applying a factor calculated from the median of all precursor intensities of each sample injection. Each precursor ion was then considered as having been identified by a tool if at least one intensity value for the 24 samples of a same dataset was reported. Each was considered quantifiable with a tool if 3 intensity values were reported in the 3 replicates in at least one of the 8 UPS1 concentrations. Missing values were then imputed by a noise value corresponding to the 1^st^ percentile of all precursor intensities for each sample injection. Precursor intensities were finally summed by stripped sequences to obtain peptide quantification and by accession number for protein quantification.

For each UPS1 concentration, the coefficient of variation over the three technical replicates was computed as CV (%) = 100 * standard deviation/mean. To evaluate linearity, only proteins corresponding to precursors having intensity values in all 3 injection replicates at 50 fmol/μg UPS1 before missing value imputation were considered. The quantification and imputed values of the 3 replicates of each concentration were then averaged and log_2_ transformed. Linear regression of log_2_(intensity) and log_10_(UPS1 concentration) was then used to obtain coefficients of determination (*r^2^*).

All 28 possible pairwise comparisons between two UPS1 concentrations were performed to assess quantification accuracy, as well as the sensitivity and specificity of each workflow (combination of DIA acquisition scheme (*Narrow, Overlapped, Mixed, Wide)*, and processing mode (*Library* or *FASTA*)) in differential expression analysis. For each comparison, only precursors having 3 quantification values before imputing missing values in one of the 2 groups being compared were summed to obtain protein quantification. Protein ratios and the corresponding log_2_ of the ratio, so-called “fold change” or FC, was calculated for each comparison and measurement accuracy was assessed by computing the mean absolute percentage error: *MAPE = 1/n * (|expected FC – experimental FC| / expected FC) * 100* (with *n* = number of UPS1 proteins quantified). Welch test *p* values were also calculated and adjusted by the Benjamini-Hochberg method for multiple testing, and the resulting *q* values were used to plot ROC curves using the pROC R package^31^. A protein was considered to differ from one sample to another (differentially expressed protein or DEP) between two groups of samples if *q* < 0.05 and |*z*| > 1.96 (*z* = (*x*-*μ*)/*σ* where *x* = log_2_ of the ratio; *μ* = average of all log_2_ ratios; *σ* = standard deviation of all log_2_ ratios). Proteins were classified as true positive (TP) if it varied coherently (UPS1), true negative (TN) if it did not vary (*E. coli*), false positive (FP) if an *E. coli* protein varied and false negative (FN) if a UPS1 protein concentration did not vary. Sensitivity (%) = TP / (TP+FN) * 100 and false discovery proportion (%) = FP / (FP+TP) * 100 were reported.

Coefficient of variation (CV), coefficient of determination (*r^2^*), MAPE and AUC were also calculated for peptides and proteins quantified by all the tools. To obtain the lists of these for each UPS1 concentration independently, the quantified precursor ions in common among all tools in each workflow (acquisition scheme and processing mode) were used. Since in most cases, the number of UPS1 peptides in common at 0.1 and 0.25 fmol/μg concentrations was zero, only UPS1 data for 1–50 fmol/μg concentrations were used for subsequent calculations.

## RESULTS AND DISCUSSION

### DIA analysis of the UPS1-E.coli proteomic standard

The complex proteomic standard used in this study reflected the difficulty of analyzing whole proteomes by *bottom-up* proteomics, especially of detecting and quantifying species present at low concentrations. This standard comprises an *E. coli* proteome background spiked with 48 human proteins (UPS1, Sigma) at 8 concentrations ranging from 0.1 fmol to 50 fmol per μg of *E. coli* protein (Fig. 1A) and digested with trypsin. Prior to DIA experiments, a spectral library was generated by DDA analysis of 48 fractions of an *E. coli* protein extract alone, trypsin-digested and fractionated by high-pH reversed-phase chromatography. A single injection of 200 fmol/μg of UPS1 in *E. coli* background was acquired in DDA mode as well. The final spectral libraries contain a total of 2 655 proteins and 20 662 peptides when generated by Skyline, 2 592 proteins and 23 574 peptides when generated by Spectronaut, allowing deep proteome coverage for subsequent DIA analysis.

The single injection of 200 fmol/μg UPS1 in *E. coli* was used to optimize the mass range and window sizes to be used in DIA experiments by plotting retention times in the LC gradient versus precursor m/z (Fig. 1B). This non-homogeneous distribution highlights the importance of adjusting DIA windows to maximize coverage of the eluted peptides. However, DIA acquisition parameters should represent a compromise between chromatographic resolution (cycle time), sensitivity (window size) and peptide sequence coverage (mass range). To determine if reducing DIA window size, and hence MS/MS spectrum complexity, could compensate for the loss of peptides when the mass range is reduced, we set the number of DIA windows constant at 75 to obtain the minimum of 10 measurements per chromatographic peak required for accurate quantification, considering that peptides elute in 25–30 s under our chromatographic conditions. Indeed, the transient length of the Orbitrap at a resolution of 15,000 is 32 ms, which results in a cycle time of 2.4 s when measuring 75 DIA windows.

The first SWATH-MS analysis described^11^ used 32 scans of 25 m/z to sequence peptides in the 400–1200 m/z range. These wide DIA windows allowed broad coverage of the mass range but at the cost of fragmenting many co-eluting peptides, resulting in highly complex chimeric MS2 spectra. By setting narrower DIA windows, more ions of the same species are accumulated prior to fragmentation, increasing analytical sensitivity, and reducing MS2 spectrum complexity. However, the mass range covered must be reduced to conserve the proper cycle time. To distribute the number of precursors equally, setting the variable isolation windows across the mass range as a compromise between sensitivity (window size) and sequenced mass range coverage, has been proposed^32^. It has also been shown that sensitivity can be increased without sacrificing the mass range selected for fragmentation, while using overlapping windows including two different cycles of 20 m/z DIA windows shifted by 10 m/z^22^.

Considering the methodologies described in the literature and based on the observed precursor distribution during DDA chromatographic run of the 200 fmol/μg UPS1 in *E. coli* sample analysis (Fig. 1B), 4 DIA window schemes were selected having different window sizes but all composed of 75 windows: m/z = 8 for *Narrow*, 15 for *Wide*, 8 and 15 for *Mixed* and 2 cycles of 8 shifted by 4 for *Overlapped*. It should be noted that *Narrow* and *Overlapped* provided 85.6% coverage of the precursors identified in DDA with the same gradient, while *Mixed* covered 98.7% and *Wide* 99.9%. These acquisition schemes allowed us to assess how the mass range selected for fragmentation and MS/MS spectrum complexity may influence DIA signal extraction and quantification.

Using 6 proprietary or open-source software tools to identify and quantify proteins, a total of 36 proteomic workflows were compared, corresponding to *Narrow*, *Wide*, *Mixed* and *Overlapped* scheme acquisitions processed either in a library-free manner using only a FASTA file, *FASTA mode,* (DIA-NN, DIA-Umpire, ScaffoldDIA, Spectronaut) or with a previously acquired DDA spectral library, *Library* mode (DIA-NN, OpenSWATH, ScaffoldDIA, Skyline, Spectronaut). To compare all the workflows fairly, precursor data were processed (filtered and normalized, missing values imputed, peptides and proteins aggregated) the same way for each DIA tool using R. Normalization was skipped when the tool had this function embedded (Spectronaut, ScaffoldDIA and DIA-NN). Skyline, DIA-Umpire and OpenSWATH generate crude quantification that need to be normalized. Moreover, the 6 tools report variable proportions of missing values that need to be imputed prior to data analysis. We applied basic methods that are widely used in our field and can be easily implemented by users with limited bioinformatics expertise: global median normalization, and imputation of missing values by the first percentile (Fig. S1). These methods have been assessed in different studies, and although they may not be the best suited, they provide acceptable results^33,34^. The precursor intensity distributions before and after normalization and the proportions of missing values are reported in supplementary figures S2 A, B and C.

It should be noted that 3 of our runs (2.5 fmol/μg UPS1 replicate 2, 5 fmol/μg UPS1 replicate 3, 25 fmol/μg UPS1 replicate 2 of the *Overlapped* dataset) display lower overall precursor intensities, due likely to a technical problem during data acquisition. Although this does affect our results, these runs were saved to test the ability of the tools to correct for such occurrences.

### Protein and peptide identification and quantification

Unlike DDA analysis for which the number protein/peptide identifications correspond to peptide-spectrum matches (PSM), and the number of protein/peptide quantifications corresponds to those having peak intensities, DIA relies on information extracted from MS/MS spectra acquired systematically. This requires defining the distinction between identified and quantified species. In our study, we chose to consider a precursor, peptide or protein as “identified” if the tool reported at least one intensity value among the 24 samples. For quantification, we considered only precursors with intensity values in all 3 replicates at each UPS1 concentration. This strategy avoids quantification with mainly imputed values. We were thus able to quantify 39.6–93.3% of identified peptides and 64.6–94.4% of identified proteins using one acquisition scheme and processing tool combination or another (Fig. S3). Of interest is that we noted greater variability in the number of peptides or proteins identified than in the number quantified, regardless of data processing pipeline. This was the case particularly when a DDA spectral library was used and suggests that certain tools identify signals from low-abundance proteins but with not enough reproducibility to make the signals quantifiable. DIA-NN in *FASTA* mode generally reported the highest number of identifiable proteins and peptides as well as the highest percentage of quantifiable species. In the *Library* mode, Spectronaut reported the highest number of peptides but Skyline the highest number of proteins as identified. The latter result is likely due to no protein FDR filtering available in Skyline. Spectronaut, Skyline and OpenSWATH report the highest number of quantifications.

Figure 2 shows the ability of the workflows to quantify proteins in a complex biological sample and to detect species present at low concentrations by reporting the number of *E. coli* and UPS1 quantified in each of the 8 concentrations. DIA acquisition scheme, software tool and processing mode (*Library* or *FASTA*) all appear to influence analytical efficiency.

**Figure 2.**
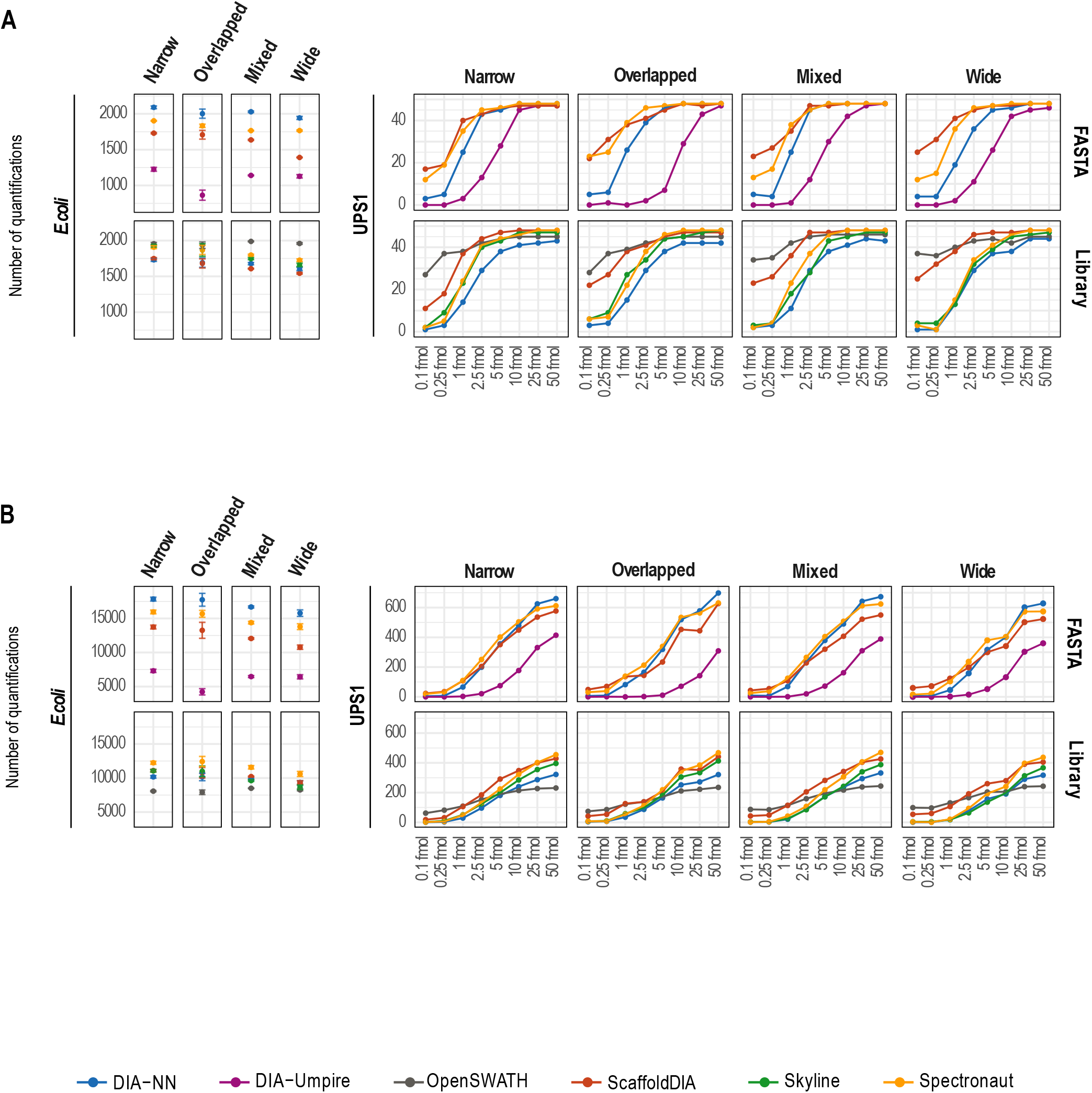
Peptide and protein quantification. The number of *E. coli* (left) or UPS1 (right) proteins (A) or peptides (B) quantified in the three replicates of each sample (*i.e.* without missing values) by the 6 software tools with each acquisition scheme (*Narrow, Overlapped, Mixed, Wide*) and data processing mode (*FASTA* or *Library*). For *E.coli*, the numbers are averaged over the 8 UPS1 concentrations. Error bars indicate the standard deviation. UPS1 numbers are averages of 3 sample replicates.

#### E. coli background

Since the *E. coli* background was the same in all samples, we averaged its peptide and protein quantifications over the 8 UPS1 concentrations for each workflow (Fig. 2A and B, left panels). All software performed slightly better with the *Narrow* window setting, for which 7.42% more proteins (on average) were quantified (Table S2A). A likely explanation is that despite its reduced precursor mass range coverage, the *Narrow* scheme yielded spectra that were less cluttered, making it easier for the algorithm to match MS2 spectra with the library or deconvolute them in library-free mode, thus improving detection. In this acquisition mode, an average of 2091 *E. coli* proteins were quantified with DIA-NN-*FASTA*, 1956 with OpenSWATH, 1926 with Skyline, 1908 with Spectronaut-*Library*, 1903 with Spectronaut-*FASTA*, 1748 with ScaffoldDIA-*Library*, 1730 with ScaffoldDIA-*FASTA*, 1731 with DIA-NN-*Library* and 1226 with DIA-Umpire. The *Narrow* and *Overlapped* window schemes allowed the highest protein sequence coverage with respectively 6.57 and 6.59 peptides per protein on average. DIA-NN-*FASTA* generally reported the highest number of quantified proteins (2016 on average for all four window schemes) and the highest sequence coverage (8.44 peptides per protein on average) followed by Spectronaut-*FASTA* (1817 proteins quantified and 8.22 peptides per protein). The numbers of proteins/peptides that all software tools were able to quantify are shown in Figure S4A. Interestingly, we calculated that only 16.5–28.8% of peptides and 39–56.8% of proteins (depending on workflow) were quantified by all tools.

#### UPS1 proteins

Figure 2 (right panels) shows that most of the tools identified at least 40 of the 48 UPS1 proteins at the 3 highest concentrations, whereas their success at the lower concentrations was variable revealing the difficulties for these tools to extract signals of low abundance species. At 1 fmol of UPS1 per μg of *E. coli* extract, ScaffoldDIA and Spectronaut in *FASTA* mode and ScaffoldDIA and OpenSWATH in *Library* mode fared better than the other workflows. At the two lowest concentrations (0.1 and 0.25 fmol/μg), these same tools still quantified more than 10 UPS1 proteins. It is interesting that the software tools reporting the highest number of proteins at low UPS1 concentration are also those having the lowest number of missing values before imputation (Fig. S2C). The number of peptides quantified dropped with concentration and the results are roughly similar for all tools except DIA-Umpire and OpenSWATH, which were less apt. Finally, the acquisition scheme did not affect the number of UPS1 proteins quantified significantly, even though *Overlapped* and *Mixed* might have been expected to perform better than the others.

### Reproducibility between technical replicates

Testing each concentration in triplicate allowed us to calculate a coefficient of variation (CV) of peptide and protein intensities to assess signal extraction reproducibility and workflow consistency (Fig. 3 and Table S2B). For *E. coli* peptides and proteins, only OpenSWATH had a CV above 15% for *Narrow*, *Wide* and *Mixed* acquisition, affirming the high reproducibility of DIA analysis for label-free quantification. Not unexpectedly, the CV was better for UPS1 proteins and peptides at high concentrations averaging respectively 3.78% and 8.35% at 50fmol/μg, versus 17.94% and 22.94% at 1 fmol/μg. In *FASTA* mode, the CV was below 20% at all concentrations above 1 fmol/μg, with *Narrow*, *Wide* and *Mixed* schemes. In *Library* mode, the CV was satisfactory at these concentrations except for OpenSWATH, which gave a CV below 20% only at the two highest concentrations. The *Overlapped* scheme dataset had technical issues during the acquisition of 3 DIA files for one replicate each at 2.5, 5 and 25 fmol/μg, which explains the higher CV obtained for *E. coli* proteins. However, it is interesting that certain tools (DIA-NN, Spectronaut and Skyline) were better than others (ScaffoldDIA, DIA-Umpire or OpenSWATH) at correcting for this, especially for UPS1 proteins. When only proteins and peptides that were quantified by all software were considered in the calculation, the CV was small for all schemes and tools, except for the *Overlapped* acquisition scheme (Fig. S4B). This suggests that the proteins and peptides shared by all the tools are also those having the highest MS signals in the analysis.

**Figure 3.**
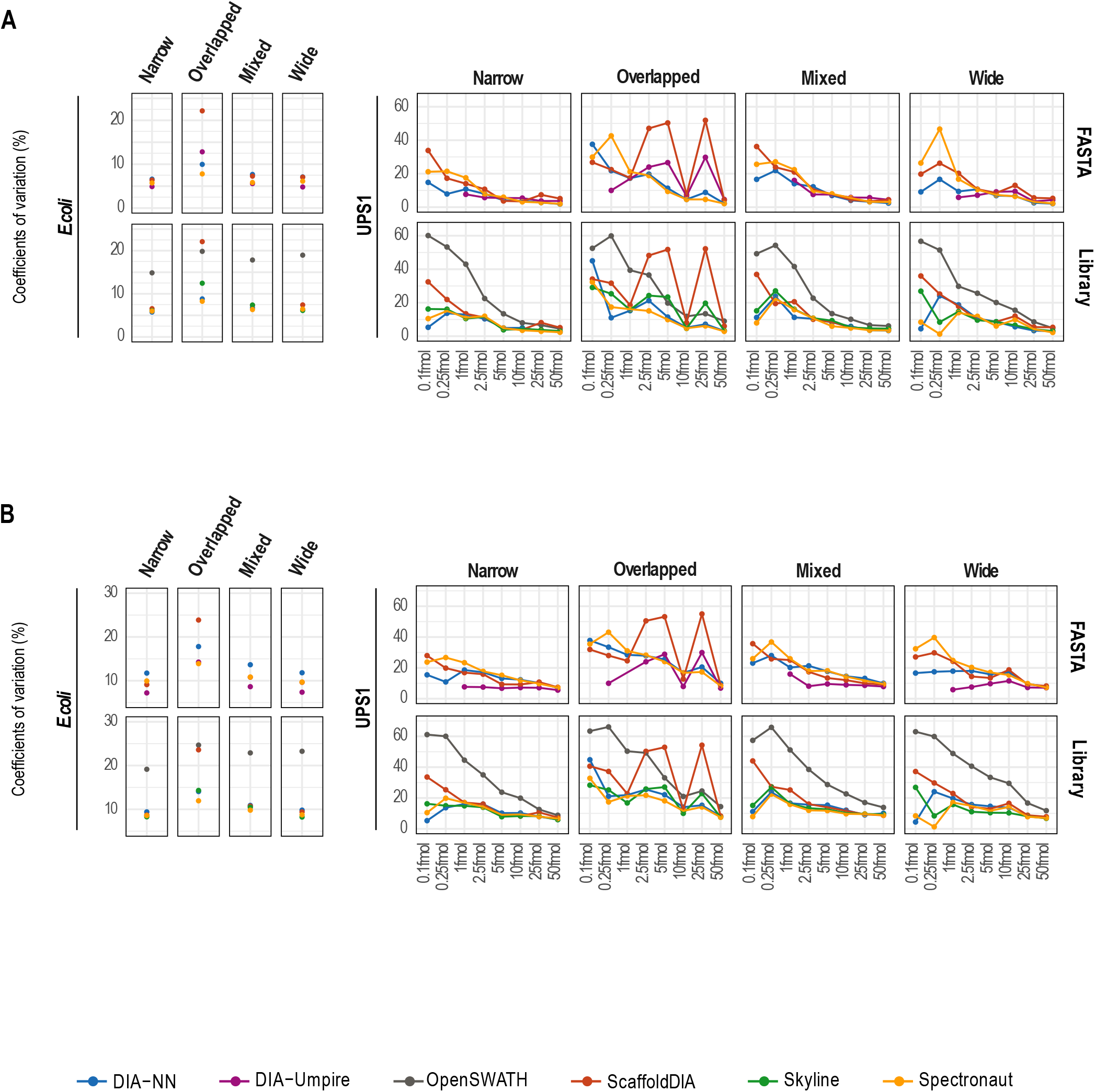
Reproducibility of quantification. Coefficient of variation of *E. coli* (left) or UPS1 (right) proteins (A) or peptides (B) based on the 3 sample replicates for the 6 software tools with each acquisition scheme (*Narrow, Overlapped, Mixed, Wide*) and data processing mode (*FASTA* or *Library*). Precursor intensities in the three replicates before missing values imputation were summed by stripped sequence to quantify peptides and by accession number to quantify proteins, for each UPS1 concentration independently. For *E. coli*, the coefficients are averaged over the 8 UPS1 concentrations.

### Linearity over the range of UPS1 protein concentrations

Another important parameter to consider when assessing the reliability of a quantification method is the linearity of the instrument response as a function of peptide concentration in the sample. To obtain accurate ratio measurements for pairwise comparisons, this response must be linear over a wide range of concentrations. Protein signal intensity distribution plotted as a function of UPS1 concentration follows the previously reported bi-linear shape^35,36^ for all workflows (Fig. S5A). It is composed of two segments: a signal segment at the highest UPS1 concentrations and a noise segment at the lowest concentrations. Since the protein intensity distributions at concentrations below 1 fmol/μg were in the noise range for most workflows, only the concentrations above this point were retained for the linear regressions, of which the coefficients of determination (*r^2^)* are shown in Fig. 4 and Table S2C. At these 6 concentrations, the *r*^2^ values were highest for DIA-NN, ScaffoldDIA, Spectronaut and Skyline, with an average above 0.96 and suggesting a very strong correlation between UPS1 concentration and protein intensity extracted by the software. Linearity was better for data acquired with the *Narrow* than with the *Wide* DIA acquisition scheme, *r^2^* averaging (all tools) respectively 0.967 and 0.957. This finding suggests again that the less cluttered MS2 spectra acquired using narrower windows made it easier to detect and quantify low-abundance signals and thus improved the overall linearity response. For narrow DIA windows, the tools able to work both in *FASTA* and *Library* modes had better *r*^2^ values in *FASTA* (0.984 versus 0.975, averaged for the three tools). Similar observations are made when only proteins quantified by all tools were considered (Fig. S4C). In addition, *r*^2^ increased for OpenSWATH and DIA-Umpire when the linear regression included only the 3, 4 or 4 highest concentrations, suggesting that more of the lower UPS1 concentrations fall in the noise segment of the bi-linear plots for these tools but that the linearity is conserved at high concentrations (Fig.S5B).

**Figure 4.**
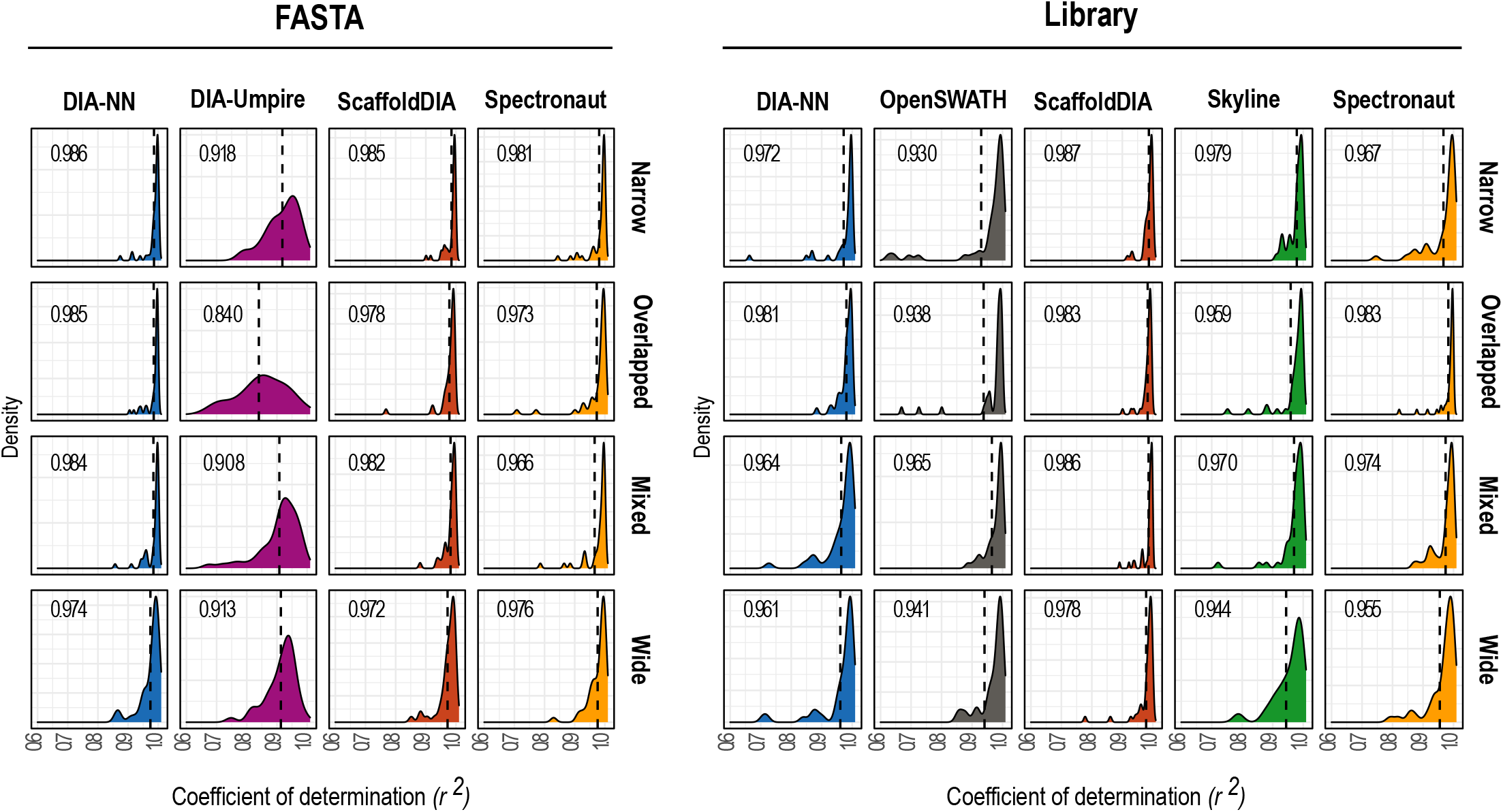
Linearity across UPS1 concentrations. Linear regression of UPS1 protein quantification over 6 concentrations (excluding the 2 lowest) in the *E. coli* protein background for each acquisition scheme (*Narrow, Overlapped, Mixed, Wide*), software tool and data processing mode (*FASTA* or *Library*). Each graph shows the coefficient of determination (*r*^2^) distribution and average (dashed vertical line and numerical value). Only proteins detected in the three replicates of the highest concentration (50 fmol/μg) before missing value imputation were included in the regression.

### Ratio accuracy in pairwise comparisons

Since the aim of most proteomics studies is to analyze differential protein expression by revealing significant quantitative differences between complex proteomes, ratio accuracy in pairwise comparisons is an important factor to consider. We therefore calculated the Fold Change (FC= log_2_(ratio) of protein intensities for each possible pair of the 8 UPS1 concentrations tested (a total of 28 pairwise comparisons) for the 36 experimental workflow conditions. The mean absolute percentage error (MAPE) to the expected FC (Fig. 5 and Table S2D) reveals an overall tendency for the error to increase when a very low UPS1 concentrations (≦ 1 fmol/μg) is compared to a high concentration. The error averaged over all workflows was 9.1% when 50 fmol and 10 fmol (expected ratio = 5) were compared, and 35% when 50 fmol and 0.1 fmol (expected ratio = 500) were compared. For comparisons of lower concentrations differing by 5-fold, (5 fmol and 1 fmol) the error averaged 33.4%. Such discrepancies in the errors might be due to the inability of the software to distinguish between signal and noise at low analyte concentrations. When both real concentrations were below 2.5 fmol/μg, the average MAPE ranged from 41.8% to 96.5%. Some workflows thus performed better than the others but nonetheless poorly. It therefore may be presumed that at 0.1 and 0.25 fmol/μg, most of the protein signals are in the noise range (Fig. S5A). However, in most studies, the protein concentration in the mixture is unknown and this error is embedded in the overall results. This weak signal recovery problem does not seem related to the DIA window width, since the MAPE did not differ significantly between *Narrow* and *Wide*. The *Library* and *FASTA* modes do not seem to be involved either, for the same reason. Based on MAPE, the workflows averaged over the ratios rank as follows (Table S2D): DIA-NN-*FASTA* (27.3%), ScaffoldDIA-*Library* (29.3%), Spectronaut-*Library* (29.4%), DIA-NN-*Library* (29.4%), Spectronaut-*FASTA* (30.9%). The more accurate tools have their own data normalization process. Although the simple normalization method that we chose (median normalization) appears to have been adequate in most cases (Fig.S2B), other methods could be applied to improve overall post-processing and correct for aberrant conditions associated with injection and acquisition^33,37^. It should be noted also that for sample concentration pairings differing by 50-fold or more, Skyline had the lowest MAPE. But its accuracy suffered at UPS1 concentrations of 10 fmol/μg or less, especially with the *Wide* acquisition scheme (which was also the case for DIA-NN). This suggests that Skyline is an excellent peak integration tool but is more easily corrupted by background noise.

**Figure 5.**
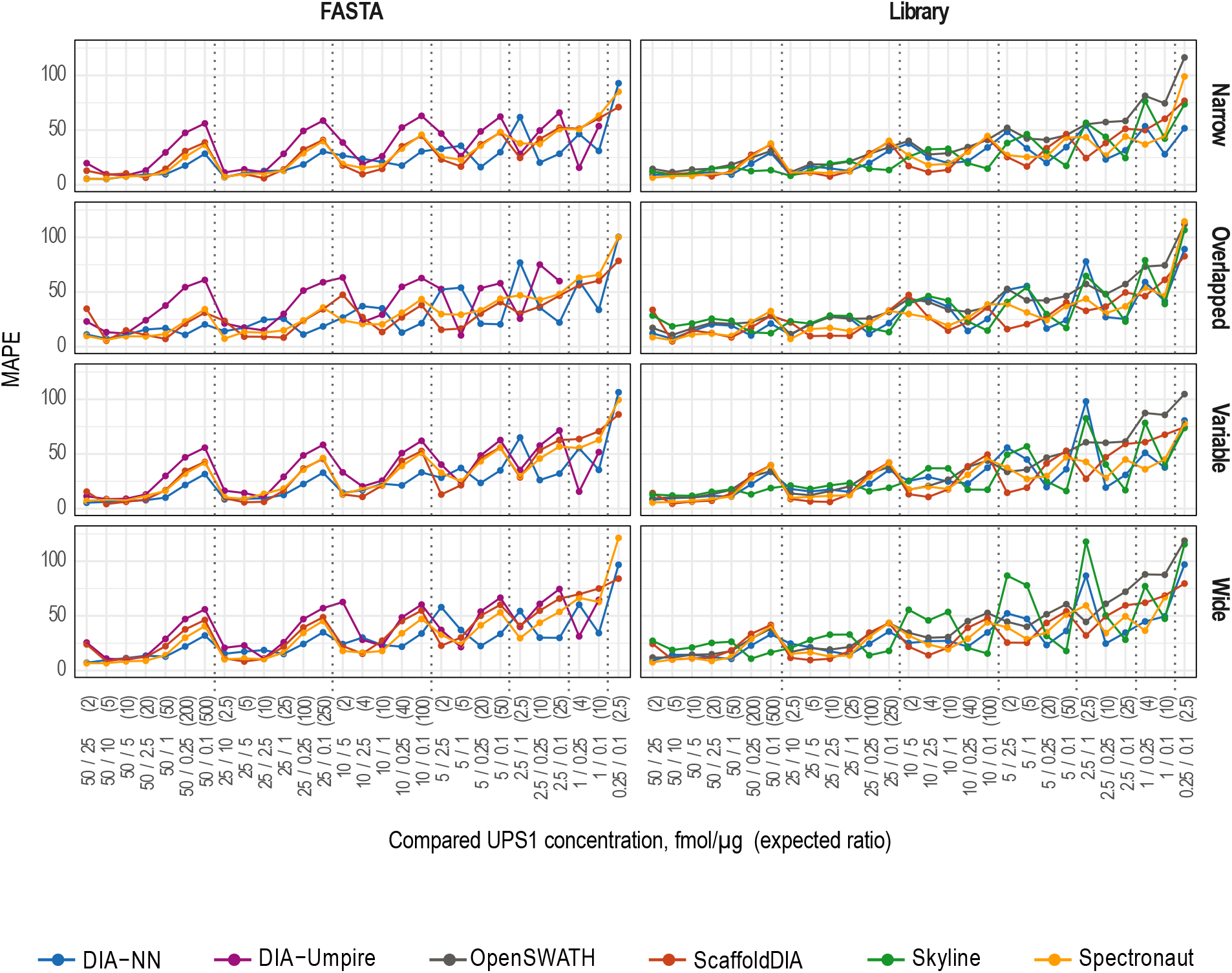
Accuracy of protein quantification. Mean absolute percentage error (MAPE) of detected UPS1 protein concentrations relative to the corresponding known concentrations for 28 paired comparisons for each acquisition scheme (*Narrow, Overlapped, Mixed, Wide*), software tool and data processing mode (*FASTA* or *Library*). *MAPE (%) = 1/n * (|Expected FC - Experimental FC| / expected FC) * 100* where FC = Log2(intensity ratio) and n= number of UPS1 protein quantified.

### Sensitivity and specificity in differential expression analysis

The use of high-performance fast-scanning LC-MS/MS instruments now allows identification and quantification of hundreds or thousands of proteins in a single analysis, as was the case in this study. However, the goal of proteome analysis is to detect differential expression of as many proteins as possible with minimal false positives by controlling the false discovery rate (FDR). Since differentially expressed proteins are known in our study (UPS1), we can assess the impact of acquisition schemes and data processing on this using receiver operating characteristic (ROC) curves of the *q* values associated with protein fold changes and by reporting the corresponding areas under the curve (AUC) (Fig. 6 and Table S2E). An AUC of 1 represents a perfect distinction between the two classes compared, namely UPS1 proteins (differentially expressed) and *E. coli* proteins (fixed background), whereas an AUC of 0.5 represents a model that is unable to distinguish the two classes. In contrast, when AUC = 0, the model is mistaking one class for the other. As expected, the software tools all seem less able to differentiate the classes at USP1 concentrations of 2.5 fmol/μg or lower. The AUC dropped from 0.962 (averaged over all workflows and comparisons) for high concentrations (at least one of the compared UPS1 concentrations ≥ 5 fmol/μg) to 0.800 for low concentrations (both ≤ 2.5 fmol/μg). This is likely due to less identification and less accurate quantification of UPS1 proteins at lower concentrations. Under these conditions, the AUC was highest using the *Narrow* windows (especially with DIA-NN, Skyline and Spectronaut) again highlighting the importance of reducing MS2 spectra complexity. We also note that the 3 aberrant MS acquisitions in the *Overlapped* dataset (due to unknown technical incidents) had an impact on AUC but not for all tools. As the reproducibility data appear to indicate (Fig. 3), Spectronaut and DIA-NN are capable of correcting for this kind of oddity and therefore report more uniform AUC values. It should also be mentioned that we used only 3 technical replicates for each sample and that better AUC values might have been obtained with more replicates, since the *q* values are influenced by this number.

**Figure 6.**
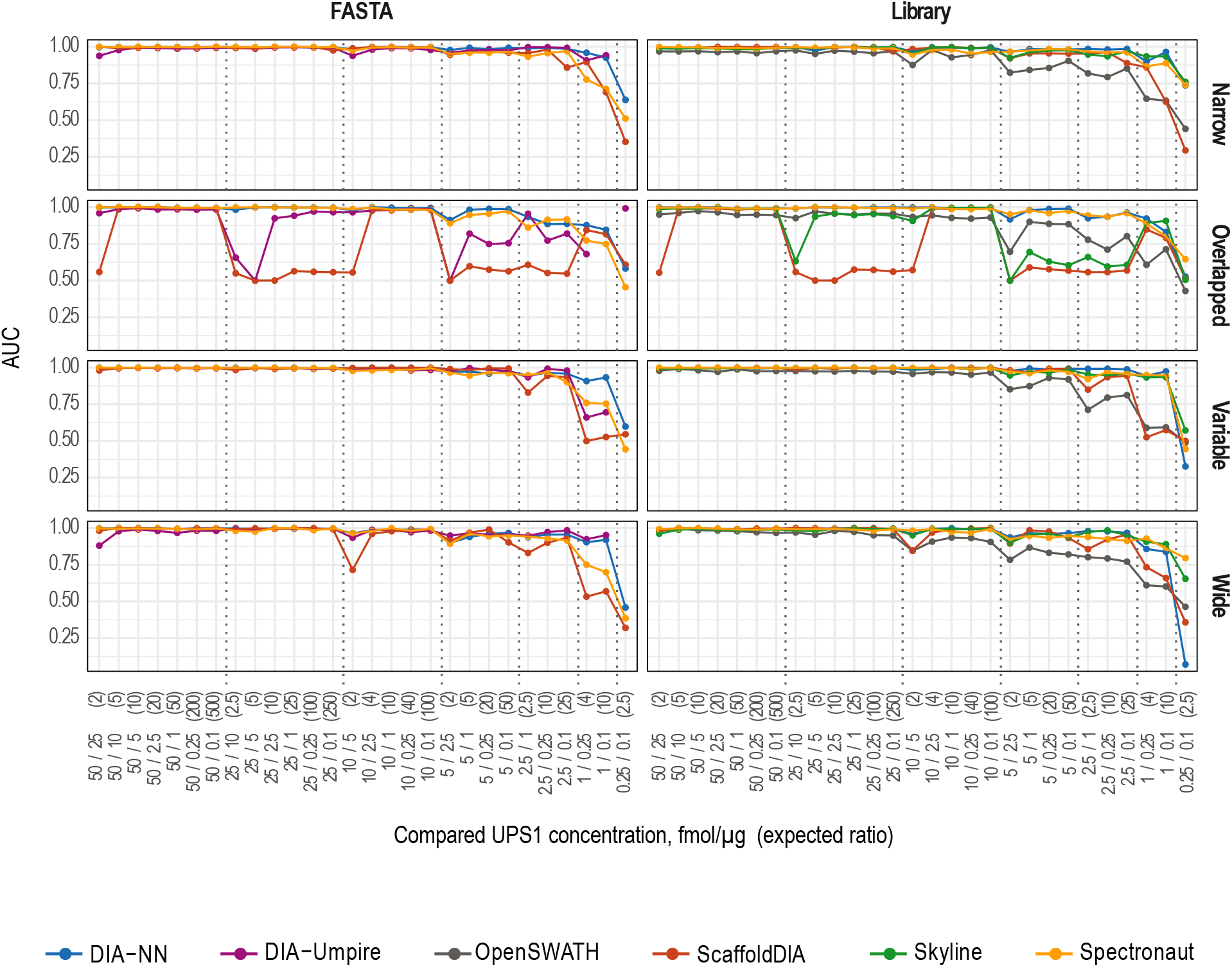
Sensitivity and specificity of the differential expression analysis. Area under the curve (AUC) corresponding to receiver operating characteristic (ROC) curves on *q* values plotted for 28 pairwise comparisons of two UPS1 concentrations for each acquisition scheme (*Narrow, Overlapped, Mixed, Wide*), software tool, and data processing mode (*FASTA* or *Library*).

Since using only *q* cut-off (often set at 0.05) to define differentially expressed protein in a proteomic analysis may lead to a high rate of false positives, in most published proteomic studies, another filtering criterion based on Fold Change or *z*-score is usually applied. The *z*-score (*z* = (FC - FC average / FC standard deviation) normalizes the fold change and thereby removes biases associated with uncentered distribution unlike when an arbitrary FC cut-off is set. To determine the adequacy of our workflows for distinguishing differential UPS1 protein expression from the background, we considered proteins as differentially expressed if they met both of the following criteria: *q* < 0.05 and |*z*| > 1.96 (corresponding to the FC distribution 95% external boundaries). We then plotted the sensitivity and false discovery proportion (FDP) obtained by counting the number of true and false positives and negatives for each of the 28 paired comparisons (Fig. S6 and Table S2F). As expected, we observed that the sensitivity of the software tool (its ability to detect differentially expressed proteins) was strongly reduced when two low UPS1 concentrations (≤ 2.5 fmol/μg) were compared, whereas certain tools (DIA-NN, Spectronaut) achieved nearly perfect performance (48 UPS1 proteins) in comparisons involving two high UPS1 concentrations, unless those concentrations differed by a small ratio (50 and 25 fmol/μg). Averaged over all tools and pairwise comparisons, the sensitivity was best with the *Narrow* (60.7%) followed by *Mixed* (53.54%) and *Wide* (49.07%) window schemes, with FDP ranging from 10.97% to 12.91%. The sensitivity was also greater in the *FASTA* mode than in the *Library* mode (53.06% versus 46.88%) with FDP of 9.52% and 10.09% respectively. Finally, the three highest sensitivities averaged over the 28 paired comparisons were obtained with DIA-NN-*Narrow-FASTA* (71.76%), Spectronaut-*Narrow-FASTA* (73.16%) and Spectronaut-*Narrow-Library* (70.20%). Among these, Spectronaut-*Narrow-FASTA* had the lowest FDP (1.88%).

### Cross-compatibility between software tools

The results presented above were computed using the standard or recommended functionalities of each software product. However, several of them accommodate spectral libraries generated using other tools. For example, Skyline and Spectronaut generate their own spectral libraries from DDA files, and these libraries are usable for quantification with DIA files in other tools. To determine whether the spectral library generation step or to the quantification step is the greater source of inconsistent results, we used our *Narrow* dataset to compare the performances of Skyline with a Spectronaut library and Spectronaut with a Skyline Library, in terms of the number of protein/peptides quantified, reproducibility (CV), linearity (*r^2^*), ratio accuracy (MAPE) and sensitivity (Fig. S7). Using the parameter settings recommended for the tool, the Spectronaut library contained 23,574 peptides versus 20,663 in the library generated by Skyline from the same DDA files. However, the number of proteins did not differ nearly as much: 2655 for Skyline and 2592 for Spectronaut (Fig. S7A). Thus, about 16% more *E.coli* peptides and 18% more UPS1 peptides were quantified using the Spectronaut library than the Skyline library whereas proteins were quantified in similar numbers (Fig. S7B). In contrast and as expected, linearity and ratio measurement accuracy seem related more to quantification performance of each tool than to the library used, with *r^2^* = 0.971 for Skyline and 0.957 for Spectronaut (Fig. S7D). As previously observed, the MAPE was lower for Skyline than Spectronaut at UPS1 concentration ratios of 40 or higher, whereas the Spectronaut MAPE was lower when both concentrations were below 10 fmol/μg (*i.e.,* middle low abundance proteins) with a smaller expected ratio (Fig. S7E).

Another example of DIA software cross-compatibility is given by DIA-Umpire and DIA-NN. These tools are able to generate a spectral library using DIA files alone without prior DDA acquisitions. Since the Quant module included in DIA-Umpire 2.0 performed poorly compared to most other tools, the developers now recommend using the DIA-Umpire module to deconvolute the DIA signal followed by a database search in MSFragger to generate the spectral library and then quantify in DIA-NN, which is also able to generate a spectral library from DIA files for this purpose. We did this using our *Narrow* dataset either with the full DIA-Umpire 2.0 pipeline, with DIA-NN in *FASTA* mode or with the spectral library generated from DIA-Umpire and MSFragger (Fig. S8). The DIA spectral library generated with DIA-NN contained 6% more peptides and 39% more proteins than that generated with DIA-Umpire. However, we note that the numbers of proteins and peptides quantified are roughly the same for both libraries when using DIA-NN for signal extraction and quantification (Fig. S8B). The comparisons of CV, *r^2^*, MAPE and AUC corroborated these results, confirming the weakness of the DIA-Umpire Quant module but validating its use for the generation of DIA spectral libraries.

### Characteristics of six DIA software tools

In addition to their performance, the choice of using a DIA software requires consideration of their cost, their user friendliness and their post-processing functionalities (Table 1). Regarding the cost, Spectronaut and ScaffoldDIA are proprietary, whereas the others are open source. The user friendliness can be determined through the presence, or not, of a graphical interface (OpenSWATH and DIA-Umpire require bioinformatics knowledge to execute the different steps in command lines) and through the ease of configuration. Some software tool configuration procedures are complicated and error prone, user manuals or reference articles may provide inadequate support and sometimes, the software development team needs to be contacted to ensure an appropriate use of the software. Therefore, the information on “ease of configuration” in Table1 represents our own experience with the different tools tested in this study. It can be different for other users depending on their experiments and knowledge of DIA methods. Another important consideration is the availability of post-processing functions: how the final data are to be reported and whether integrated intensity normalization is included. If not, presentation of the output might require bioinformatics knowledge (e.g., R language) to handle large datasets. Finally, the processing time of the software tools is also important for large-scale studies. However, the tools used in this study have different system requirements, therefore we were not able to run them in a similar environment allowing a fair comparison of their performances. Nevertheless, we observed, using software designed to accommodate both *Library* and *FASTA* analyses (*DIA-NN, ScaffoldDIA* and *Spectronaut*), that the analysis performed with a spectral library was always faster than with a *FASTA* file which could be an advantage while running large experiments.

**Table 1.**
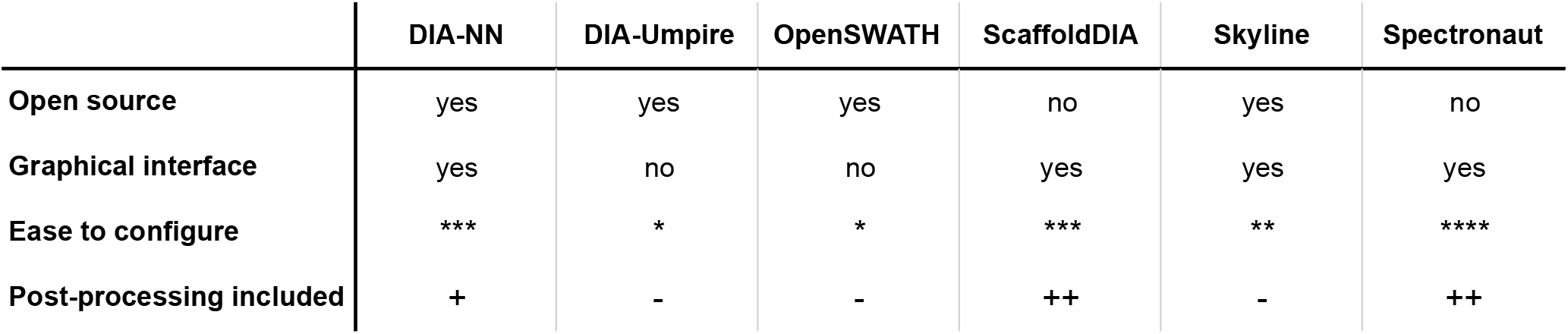
Characteristics of six DIA software tools. (****) methods/parameters are intuitive and/or well described in the documentation; (***) methods/parameters are described in the documentation but assistance (from the developer/software team) might be required to avoid error; (**) the documentation needs to be improved and/or assistance is required for first utilization(s); (*) no documentation is provided and/or bioinformatics skills are required; (−) no post-processing is offered; (+) only normalization is performed automatically, precursor data only are reported; (++) normalization is included, peptide and proteins reports are included as well as ratios and *p* values.

|  | DIA-NN | DIA-Umpire | OpenSWATH | ScaffoldDIA | Skyline | Spectronaut |
| --- | --- | --- | --- | --- | --- | --- |
| **Open source** | yes | yes | yes | no | yes | no |
| **Graphical interface** | yes | no | no | yes | yes | yes |
| **Ease to configure** | *** | * | * | *** | ** | **** |
| **Post-processing included** | + | - | - | ++ | - | ++ |

## CONCLUSIONS

In this study, we were able to assess the performance and reliability of several data-independent acquisition workflows for protein identification and quantification by processing thousands of measurements. Although certain software tools generally performed better than others under our conditions, each showed its own strengths and weaknesses in terms of protein identification and quantification, linearity, reproducibility and ratio accuracy.

The use of four different acquisition schemes showed that the size of precursor windows is an important parameter to consider when performing DIA analyses. Narrow windows generally supported better system performance than did wide windows, despite a lower coverage of the mass range. This suggests that faster scanning on the most recent Orbitrap instruments in conjunction with the use of a larger mass range and/or reduced window width should contribute to improving DIA analysis performance. Our results also suggest that some published acquisition schemes (overlapped or mixed windows) do not perform significantly better than conventional settings when analyzing complex mixtures of proteins in our conditions, nor does the use of a very large spectral library acquired from 48 fractions of *E. coli* improve the number of proteins identified and quantified. Indeed, the signals of low-abundance peptides detected by this extensive fractionation are not easily recovered by DIA processing tools fed unfractionated DIA analyses. This finding corroborates recent studies^38,39^ that question the usefulness of libraries generated prior to DIA analysis. However, we observed that post-acquisition analysis was always faster with a spectral library than with a FASTA file, which would be advantageous for large experiments. Other library types such as repetitive measurement of non-fractionated sample library^39^ or chromatogram library^20^ might improve the results shown here. Furthermore, publicly available libraries^40^, gas-phase fractionation^41^ or *in silico* generated libraries^42^ could also be used to avoid multiple injections per sample.

This study shows that the choice of processing software is the greatest source of quality issues affecting DIA-based proteomics analyses, followed by the size and number of DIA windows and finally the decision to use or not a spectral library. In addition to performance, cost, user friendliness and post-processing functionalities must be considered. Finally, processing time is an important consideration for large-scale studies. Our only basis for evaluating this aspect was the comparison of FASTA and Libraries, since the tools used in this study were not used in a similar environment.

Our analysis of 96 DIA raw files from 36 workflows and evaluation of quantification is so far the most comprehensive comparison available of DIA methods combined with Orbitrap mass spectrometers, the most widely used instruments in the proteomic research community. These results should help scientists choose the acquisition parameters and software processing that are best suited to their DIA application, and with all our data being available on ProteomeXchange repository, other teams will be able to reapply their own statistics or test other software tools. Moreover, it may help developers to improve their algorithms and therefore increase the capabilities of the DIA strategy in proteomics studies.

## ASSOCIATED CONTENT

### Supporting Information

The Supporting Information is available free of charge at:

Figure S1. Data processing. Figure S2. Normalization and missing values information. Figure S3. Peptides and proteins identified and quantified. Figure S4. Benchmark of DIA workflows considering only analytes that all tools detected. Figure S5. Evaluation of data linearity. Figure S6. Differential analysis. Figure S7. Software cross-compatibility - Quantification results using Skyline and Spectronaut. Figure S8. Software cross-compatibility - Quantification results using DIA-Umpire and DIA-NN (PDF)

Table S1. Acquisition and processing parameter settings (XLSX)

Table S2. Quantification data summary (XLSX)

## AUTHOR INFORMATION

### Author contributions

C.G and F.R.D. designed experiments; C.G., C.J.B., M.L. and L.M. performed experiments; C.G. and F.R.D. performed data analysis; C.G and F.R.D. wrote the manuscript; A.D. supervised the research.

### Notes

The authors declare no competing interests.

All raw mass spectrometry data, software outputs and quantification results are publicly available on ProteomeXchange repository (www.proteomexchange.org) with the identifier PXD026600. R scripts used to compute the data are available on GitHub: https://github.com/ArnaudDroitLab/DIA_Paper

## ACKNOWLEDGMENTS

The authors are grateful to David Bouyssié for critical reading of the manuscript.

