## Supplementary Figures S1-S8 for "Extensive and accurate benchmarking of DIA acquisition methods and software tools using a complex proteomic standard"

### Table of contents

#### List of figures


**Supplementary Figure S1** - Data processing **Page S-3**

**Supplementary Figure S2** - Normalization and missing values information **Page S-4**

**Supplementary Figure S3** - Peptides and proteins identified and quantified **Page S-7**

**Supplementary Figure S4** - Benchmark of DIA workflows considering only analytes that all tools detected **Page S-8**

**Supplementary Figure S5** - Evaluation of data linearity **Page S-10**

**Supplementary Figure S6** - Differential analysis **Page S-12**

**Supplementary Figure S7** - Software cross-compatibility - Quantification results using Skyline and Spectronaut **Page S-15**

**Supplementary Figure S8** - Software cross-compatibility - Quantification results using DIA-Umpire and DIA-NN **Page S-16**


#### List of tables


**Supplementary Table S1** - Acquisition and processing parameter settings

**Supplementary Table S2** - Quantification data summary

**Supplementary Figure 1**

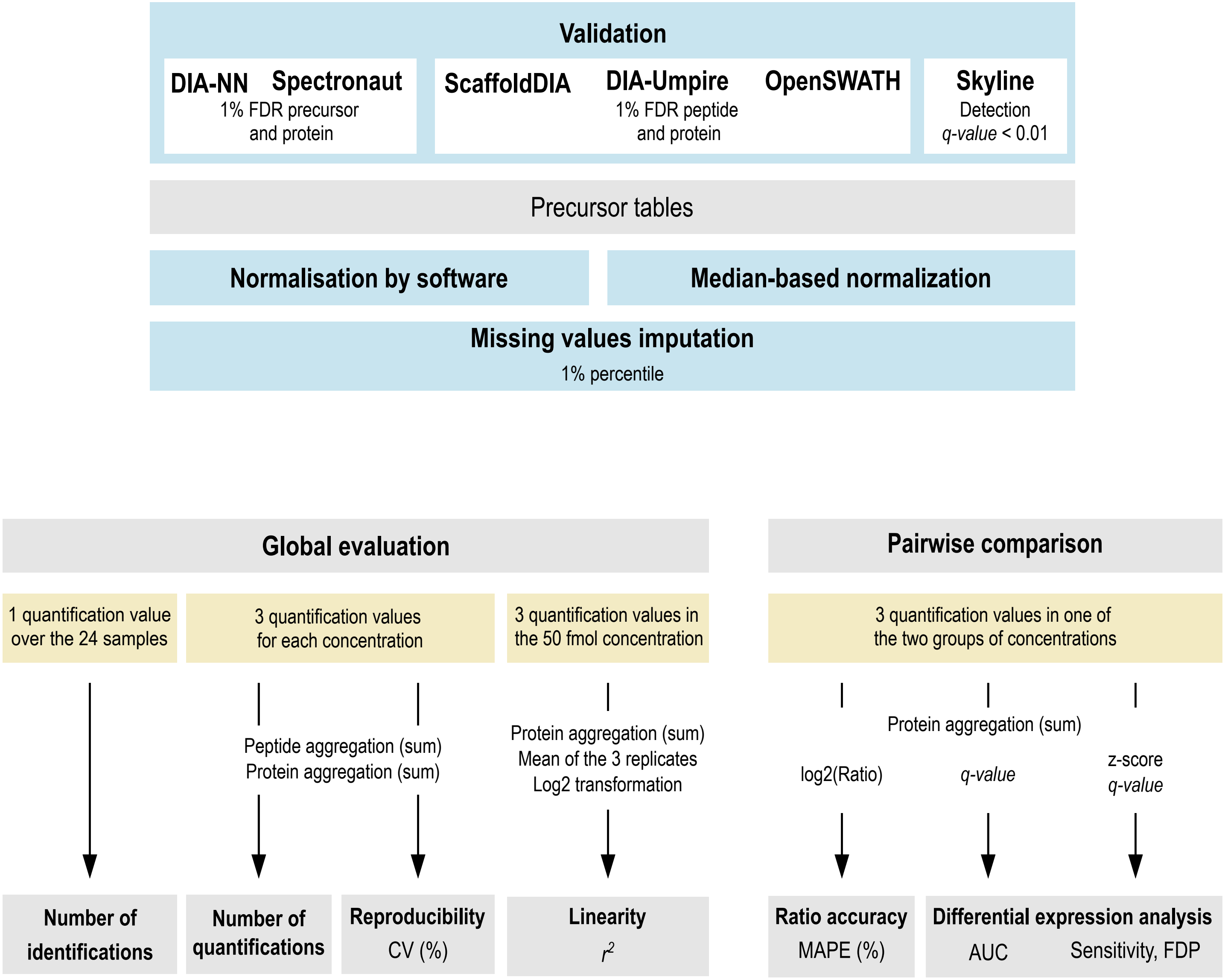


**Data processing:** Post-processing (validation, normalization, missing values imputation) (*blue boxes*) applied to DIA software outputs; Filtering (*yellow boxes*) applied to obtain: (*left*) the number of peptides or proteins identified and quantified, reproducibility and linearity data in a global evaluation and (*right*) the protein ratio accuracy, sensitivity, false discovery proportion in pairwise comparisons.

Supplementary Figure 2

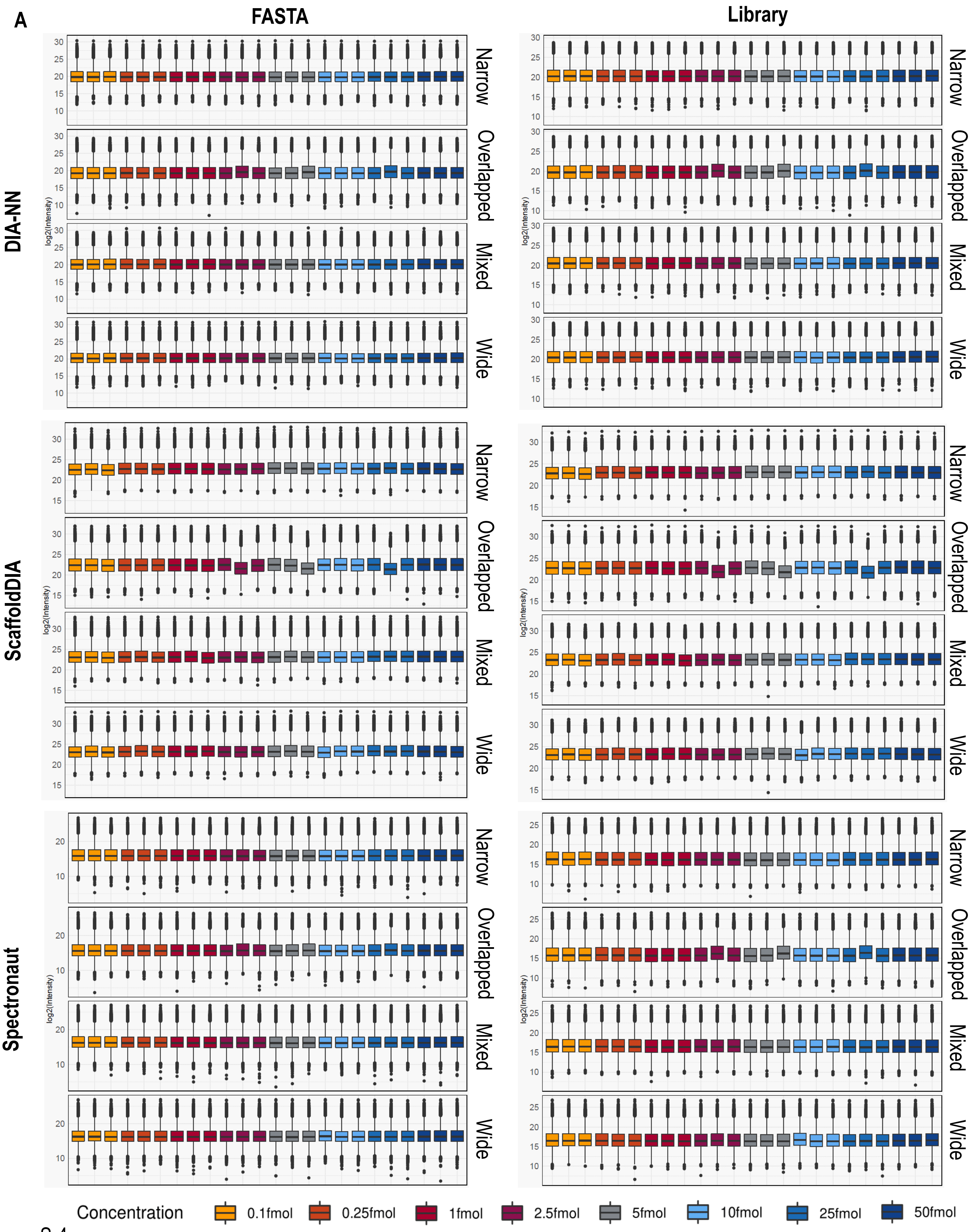

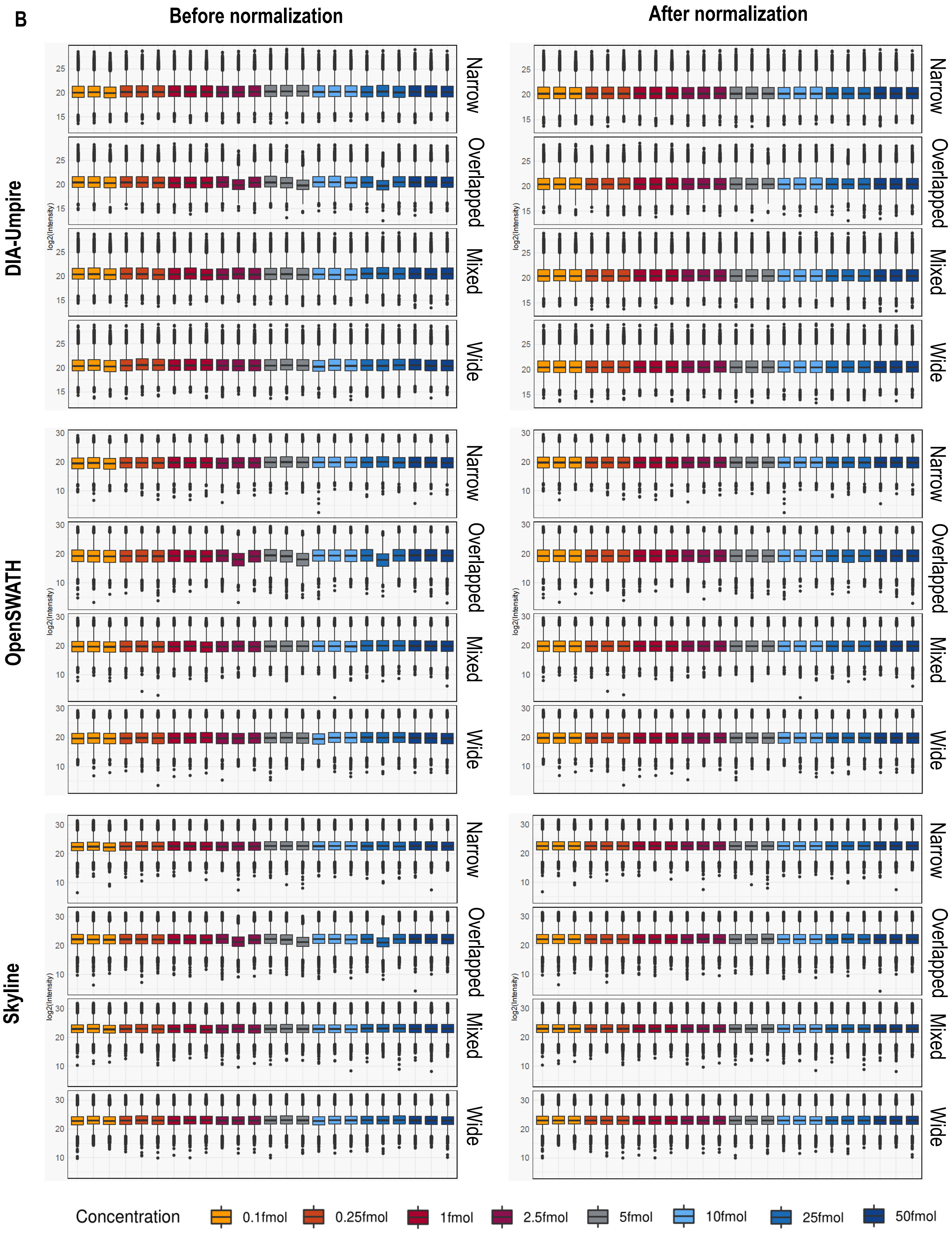
B
Before normalization
After normalization
DIA-Umpire
OpenSWATH
Skyline
log2(Intensity)
Narrow
Overlapped
Mixed
Wide
Concentration
0.1fmol
0.25fmol
1fmol
2.5fmol
5fmol
10fmol
25fmol
50fmol

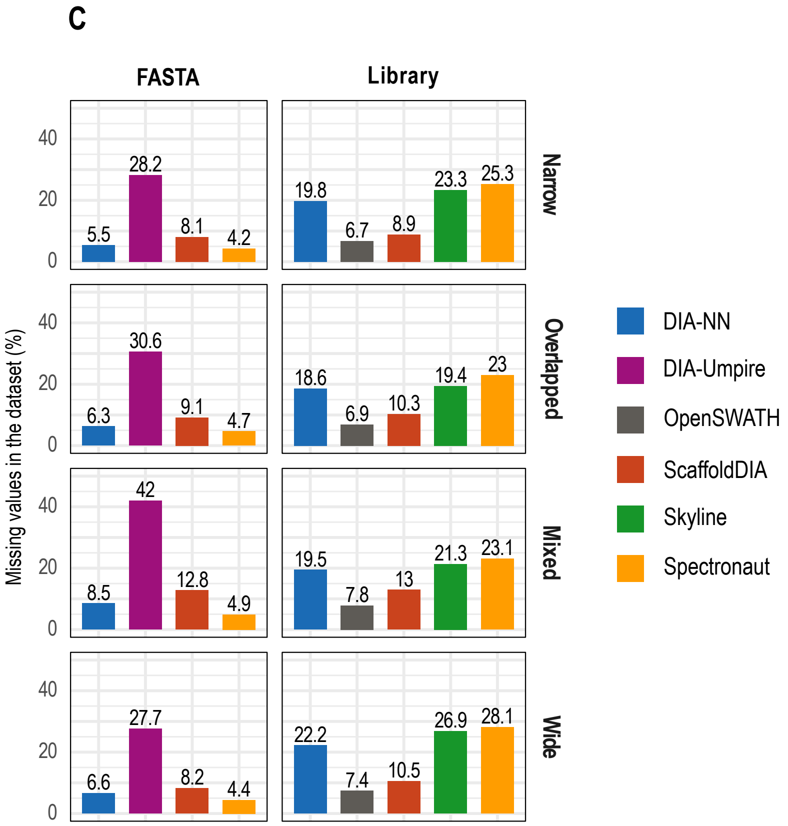


**Normalization and missing values** - (**A**) Distribution of $log_2$ intensities obtained using DIA-NN, ScaffoldDIA and Spectronaut with automatic normalization by software, in *FASTA* or *Library* mode; (**B**) Distribution of $log_2$ intensities obtained using DIA-Umpire, OpenSWATH and Skyline before and after median-based normalization; (**C**) Percentage of missing values reported for each dataset before imputation.

**Supplementary Figure 3**

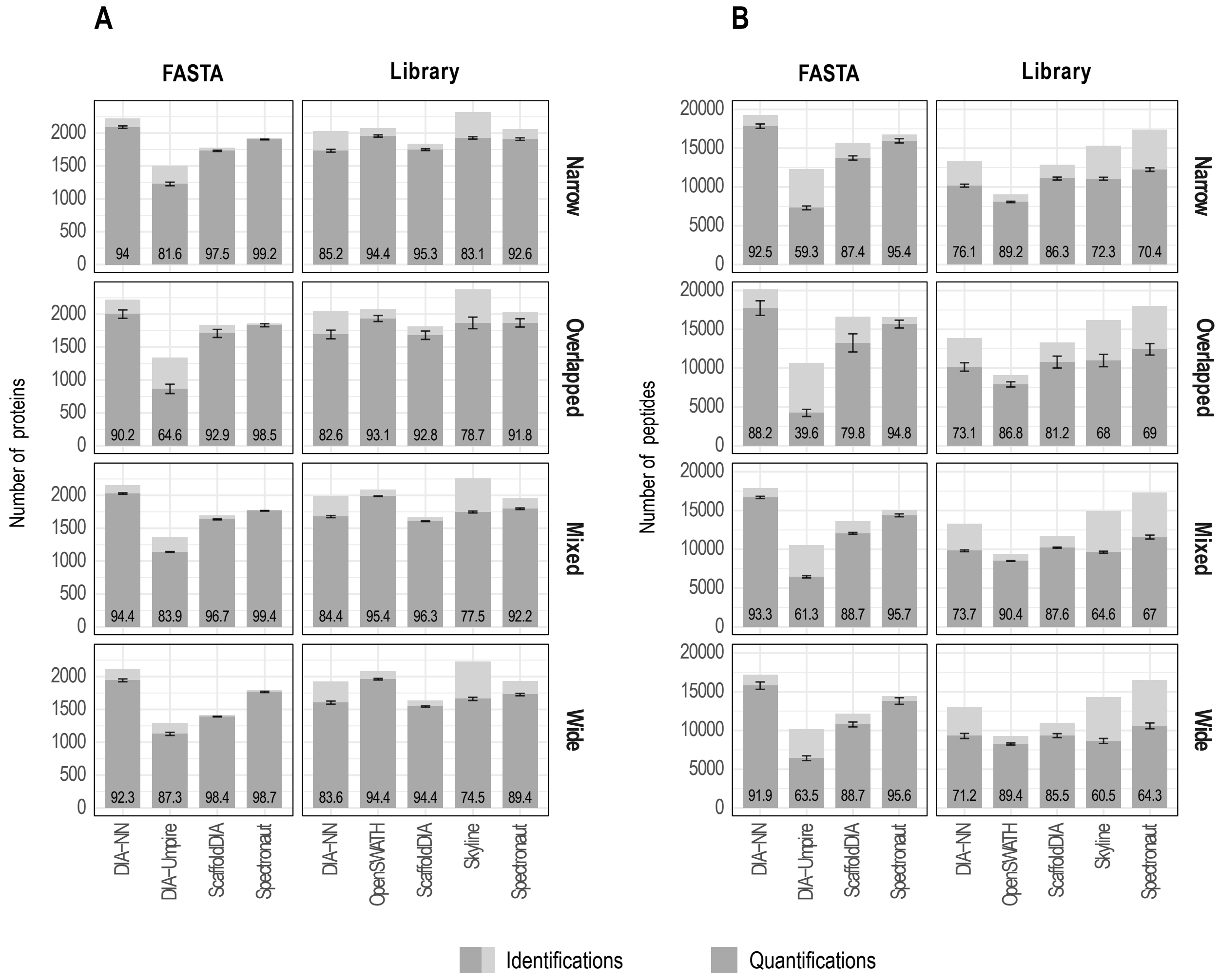


**Peptides and proteins identified and quantified** - Number of *E. coli* and UPS1 proteins (**A**) or peptides (**B**) identified (sum of *light* and *dark grey*) or quantified (*dark grey* only). To qualify as 'identified' a peptide or protein intensity must be measured in at least one of the 24 samples; a peptide or protein was 'quantified' in one condition (UPS1 concentration) if an intensity was measured in the 3 replicates at this at each UPS1 concentration, the average over the 8 UPS1 concentrations is reported. Error bars indicate standard deviation. The number at the bottom of each bar is the % of the total number of proteins identified that were also quantified.

### Supplementary Figure 4

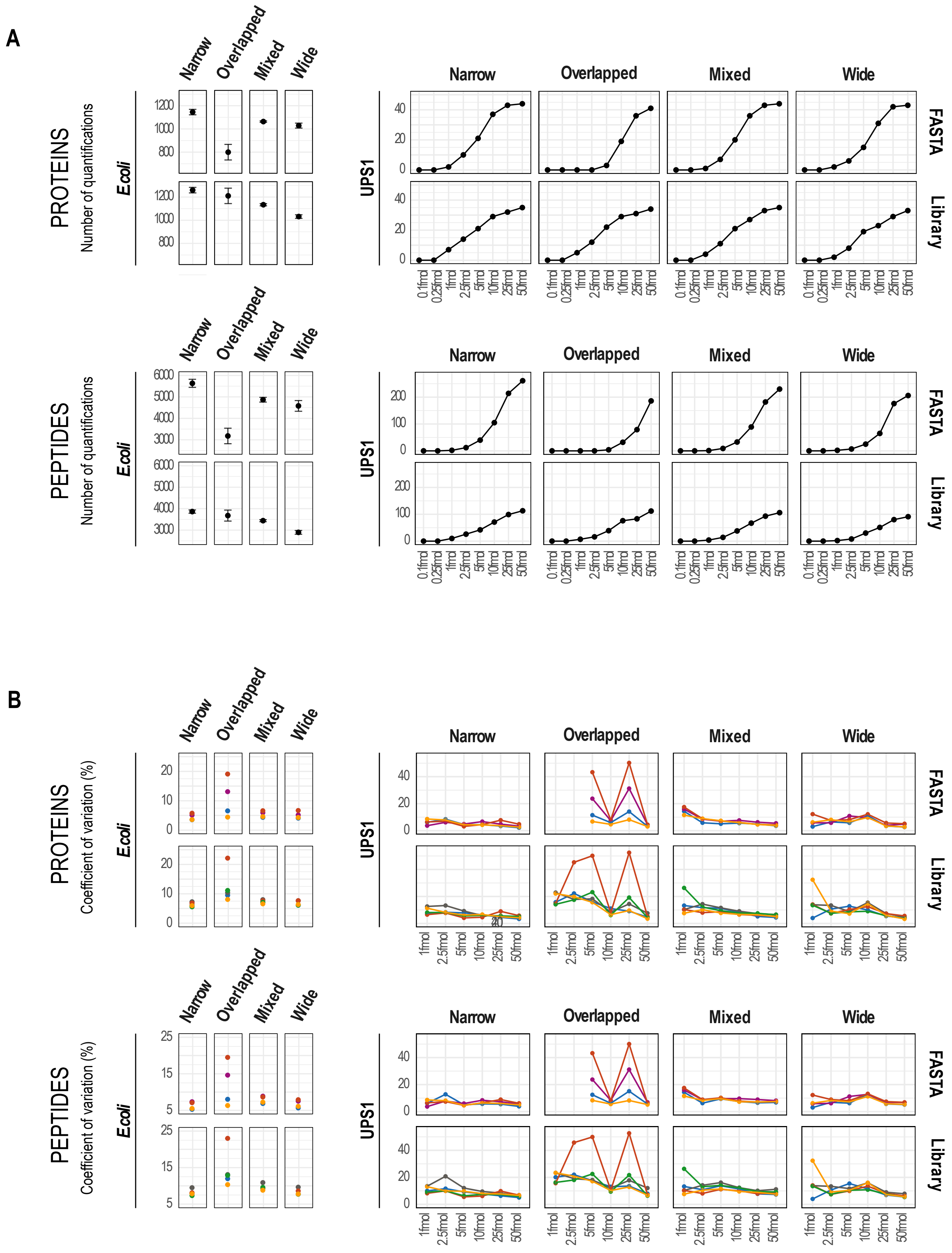

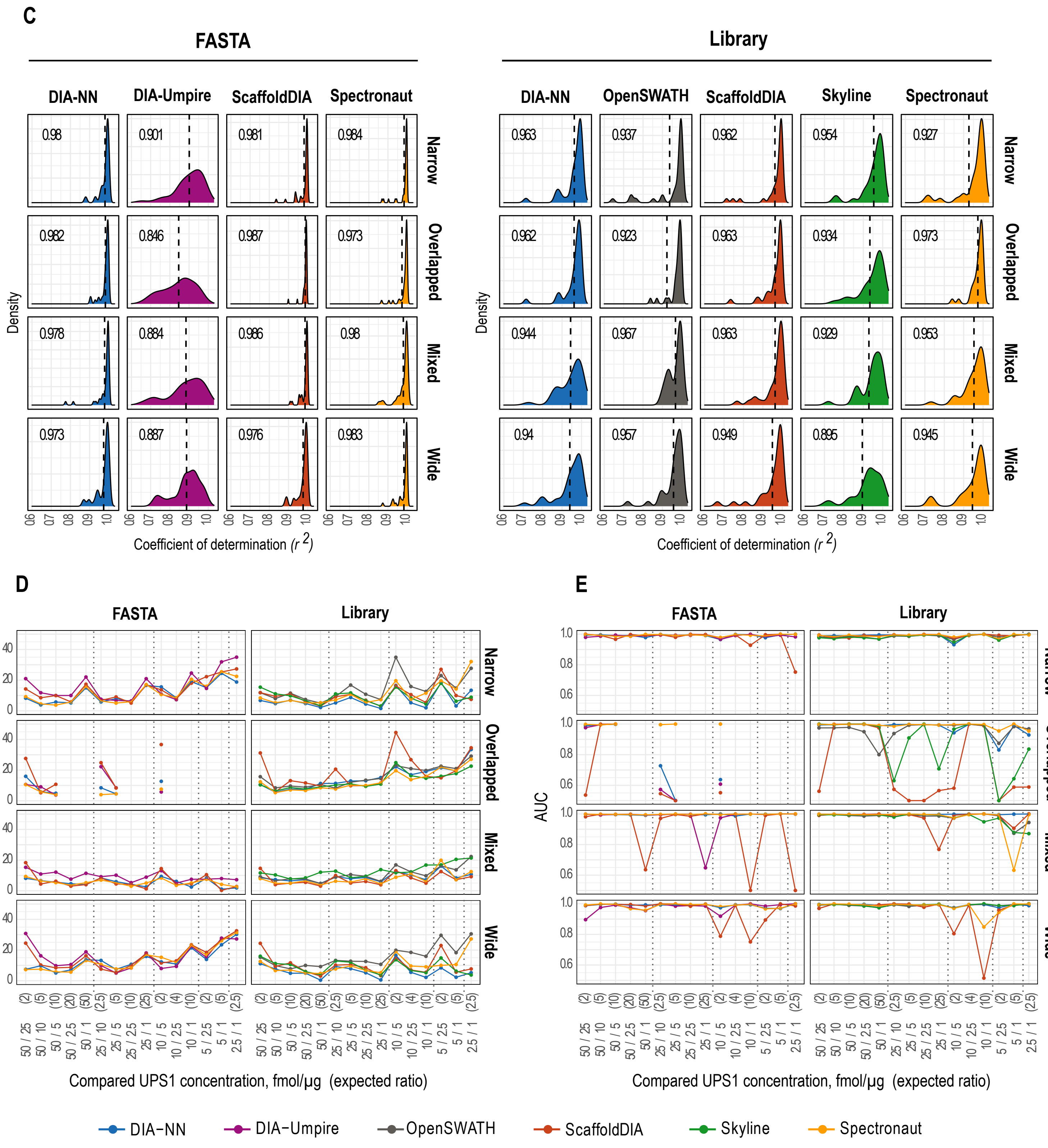


**Benchmark of DIA workflows considering only analytes that all tools detected** - For each workflow (acquisition scheme (*Narrow, Overlapped, Mixed, Wide*) and processing mode (*FASTA* or *Library*)) and for each of the 8 concentrations, only precursors quantified by all the tools were used to report: (**A**) Number of peptides and proteins quantified; (**B**) Coefficient of variation (%); (**C**) UPS1 coefficient of determination ($r^2$); (**D**) Mean absolute percentage error (MAPE) of the Fold Change in pairwise comparisons and (**E**) Area under the curve (AUC) from ROC curves.

Supplementary Figure 5

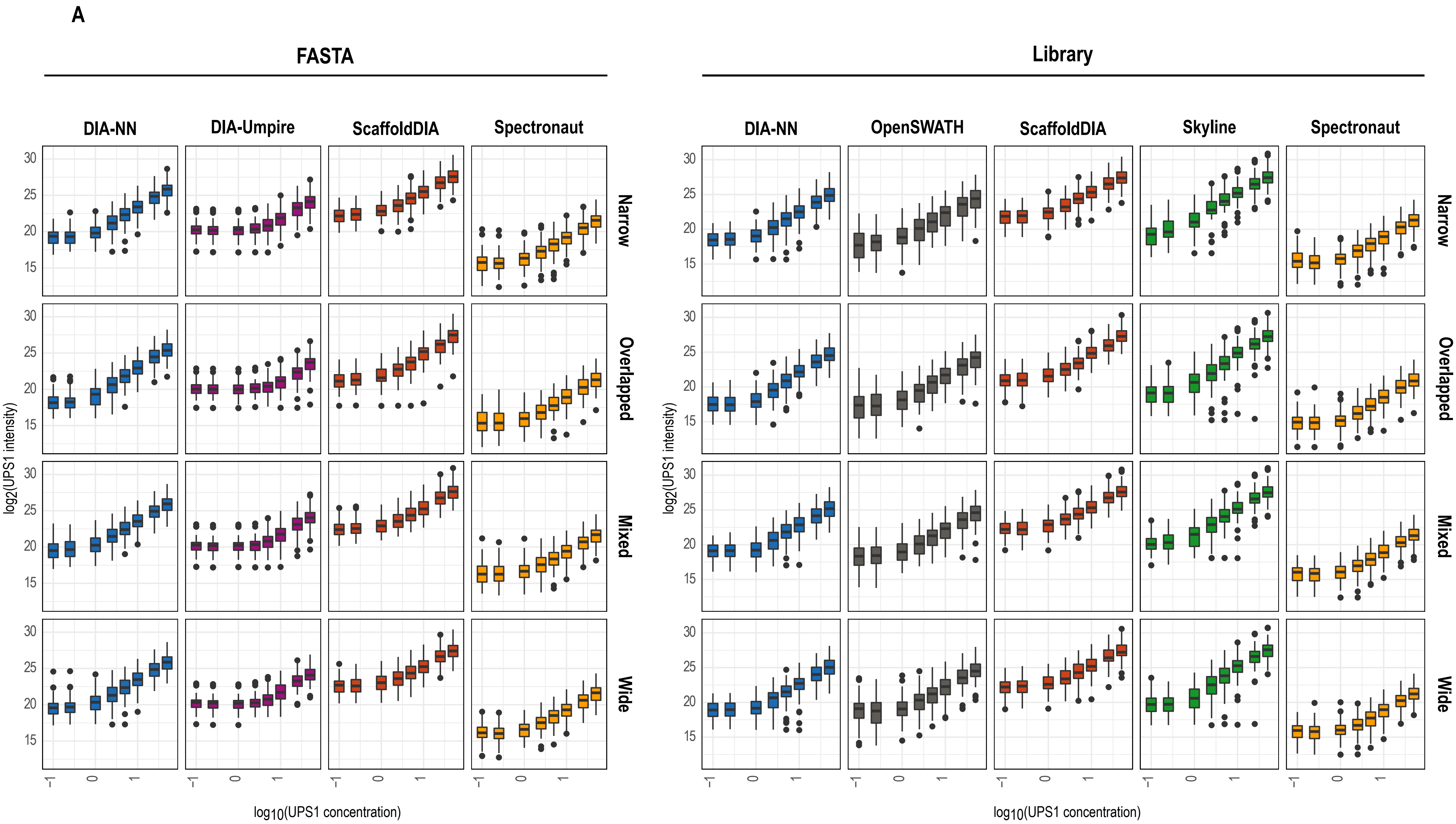

**B**

| | | | 0.1fmol to 50fmol | 0.25fmol to 50fmol | 1fmol to 50fmol | 2.5fmol to 50fmol | 5fmol to 50fmol | 10fmol to 50fmol |
|---|---|---|---|---|---|---|---|---|
| FASTA | DIA-NN | Narrow | 0.909 | 0.946 | 0.986 | 0.993 | 0.993 | 0.999 |
| FASTA | DIA-NN | Overlapped | 0.920 | 0.953 | 0.985 | 0.991 | 0.996 | 0.996 |
| FASTA | DIA-NN | Mixed | 0.904 | 0.940 | 0.984 | 0.994 | 0.997 | 0.999 |
| FASTA | DIA-NN | Wide | 0.889 | 0.924 | 0.974 | 0.982 | 0.991 | 0.997 |
| FASTA | DIA-Umpire | Narrow | 0.723 | 0.802 | 0.918 | 0.974 | 0.990 | 0.994 |
| FASTA | DIA-Umpire | Overlapped | 0.655 | 0.726 | 0.840 | 0.896 | 0.944 | 0.980 |
| FASTA | DIA-Umpire | Mixed | 0.721 | 0.799 | 0.908 | 0.970 | 0.989 | 0.995 |
| FASTA | DIA-Umpire | Wide | 0.717 | 0.799 | 0.913 | 0.965 | 0.975 | 0.987 |
| FASTA | ScaffoldDIA | Narrow | 0.906 | 0.936 | 0.985 | 0.995 | 0.997 | 0.999 |
| FASTA | ScaffoldDIA | Overlapped | 0.886 | 0.924 | 0.978 | 0.989 | 0.985 | 0.976 |
| FASTA | ScaffoldDIA | Mixed | 0.883 | 0.922 | 0.982 | 0.995 | 0.997 | 0.995 |
| FASTA | ScaffoldDIA | Wide | 0.873 | 0.916 | 0.972 | 0.991 | 0.991 | 0.990 |
| FASTA | Spectronaut | Narrow | 0.881 | 0.929 | 0.981 | 0.993 | 0.996 | 1.000 |
| FASTA | Spectronaut | Overlapped | 0.869 | 0.916 | 0.973 | 0.993 | 0.996 | 0.999 |
| FASTA | Spectronaut | Mixed | 0.865 | 0.913 | 0.966 | 0.982 | 0.998 | 0.999 |
| FASTA | Spectronaut | Wide | 0.871 | 0.920 | 0.976 | 0.983 | 0.986 | 0.994 |
| Library | DIA-NN | Narrow | 0.881 | 0.917 | 0.972 | 0.982 | 0.984 | 0.999 |
| Library | DIA-NN | Overlapped | 0.895 | 0.939 | 0.981 | 0.989 | 0.993 | 0.997 |
| Library | DIA-NN | Mixed | 0.860 | 0.907 | 0.964 | 0.982 | 0.994 | 0.997 |
| Library | DIA-NN | Wide | 0.848 | 0.895 | 0.961 | 0.978 | 0.989 | 0.994 |
| Library | OpenSWATH | Narrow | 0.858 | 0.880 | 0.930 | 0.952 | 0.956 | 0.995 |
| Library | OpenSWATH | Overlapped | 0.853 | 0.895 | 0.938 | 0.958 | 0.973 | 0.970 |
| Library | OpenSWATH | Mixed | 0.859 | 0.903 | 0.965 | 0.973 | 0.987 | 0.993 |
| Library | OpenSWATH | Wide | 0.803 | 0.868 | 0.941 | 0.966 | 0.976 | 0.992 |
| Library | ScaffoldDIA | Narrow | 0.906 | 0.943 | 0.987 | 0.994 | 0.996 | 0.999 |
| Library | ScaffoldDIA | Overlapped | 0.900 | 0.942 | 0.983 | 0.990 | 0.985 | 0.976 |
| Library | ScaffoldDIA | Mixed | 0.893 | 0.934 | 0.986 | 0.996 | 0.997 | 0.996 |
| Library | ScaffoldDIA | Wide | 0.892 | 0.927 | 0.978 | 0.990 | 0.992 | 0.991 |
| Library | Skyline | Narrow | 0.947 | 0.952 | 0.979 | 0.987 | 0.993 | 0.998 |
| Library | Skyline | Overlapped | 0.906 | 0.935 | 0.959 | 0.974 | 0.975 | 0.983 |
| Library | Skyline | Mixed | 0.912 | 0.938 | 0.970 | 0.982 | 0.992 | 0.995 |
| Library | Skyline | Wide | 0.882 | 0.920 | 0.944 | 0.958 | 0.975 | 0.989 |
| Library | Spectronaut | Narrow | 0.841 | 0.904 | 0.967 | 0.985 | 0.992 | 0.998 |
| Library | Spectronaut | Overlapped | 0.869 | 0.915 | 0.983 | 0.986 | 0.994 | 0.998 |
| Library | Spectronaut | Mixed | 0.846 | 0.906 | 0.974 | 0.990 | 0.998 | 0.999 |
| Library | Spectronaut | Wide | 0.821 | 0.881 | 0.955 | 0.985 | 0.993 | 0.997 |

Color scale: 0.6, 0.7, 0.8, 0.9, 1.0

**Evaluation of data linearity** - For each workflow (**A**) Boxplots of UPS1 protein intensity distribution at each UPS1 concentration; (**B**) Average coefficient of determination ($r^2$) of linear regressions calculated using 3 to 6 UPS1 concentrations.

### Supplementary Figure S6

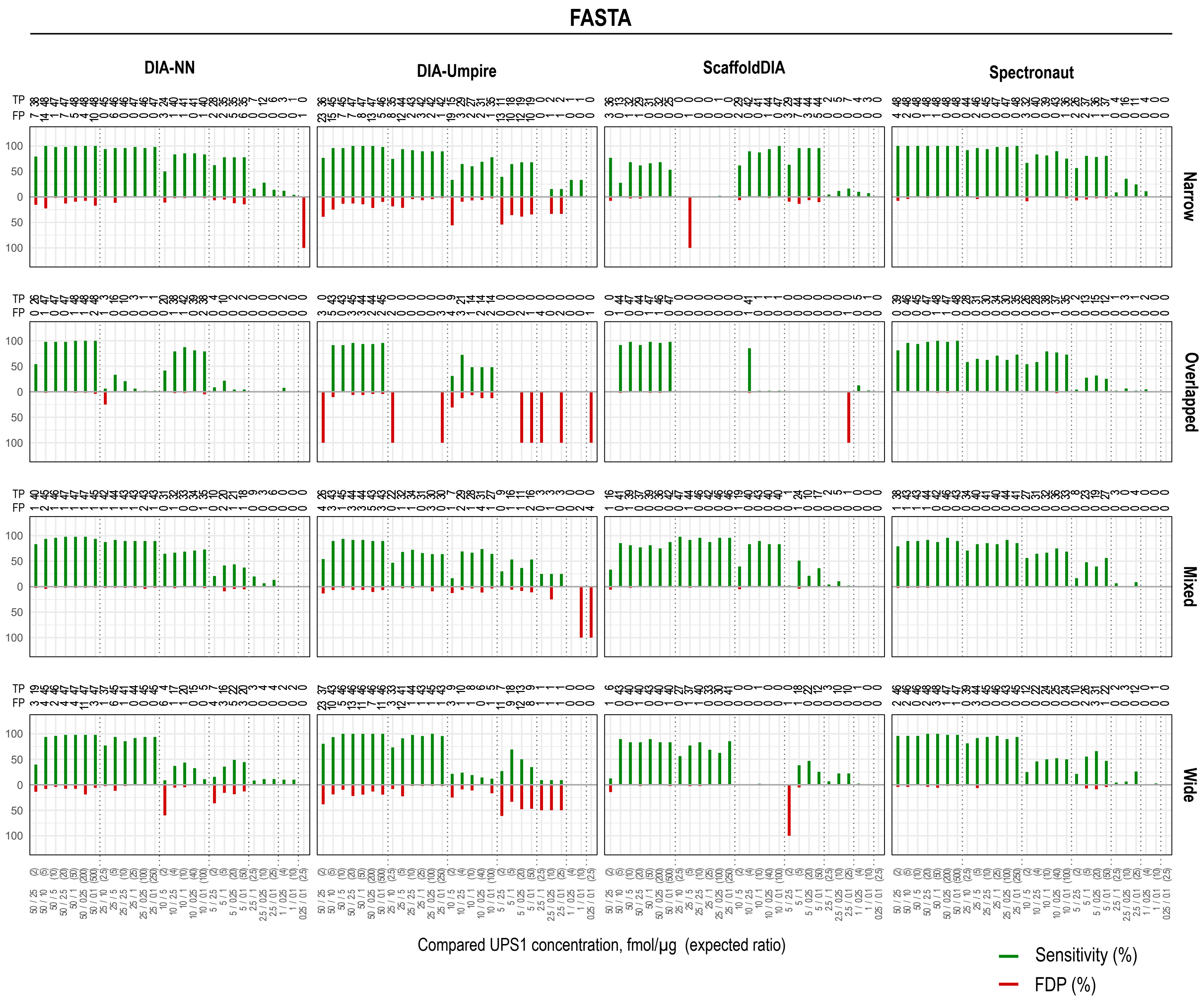

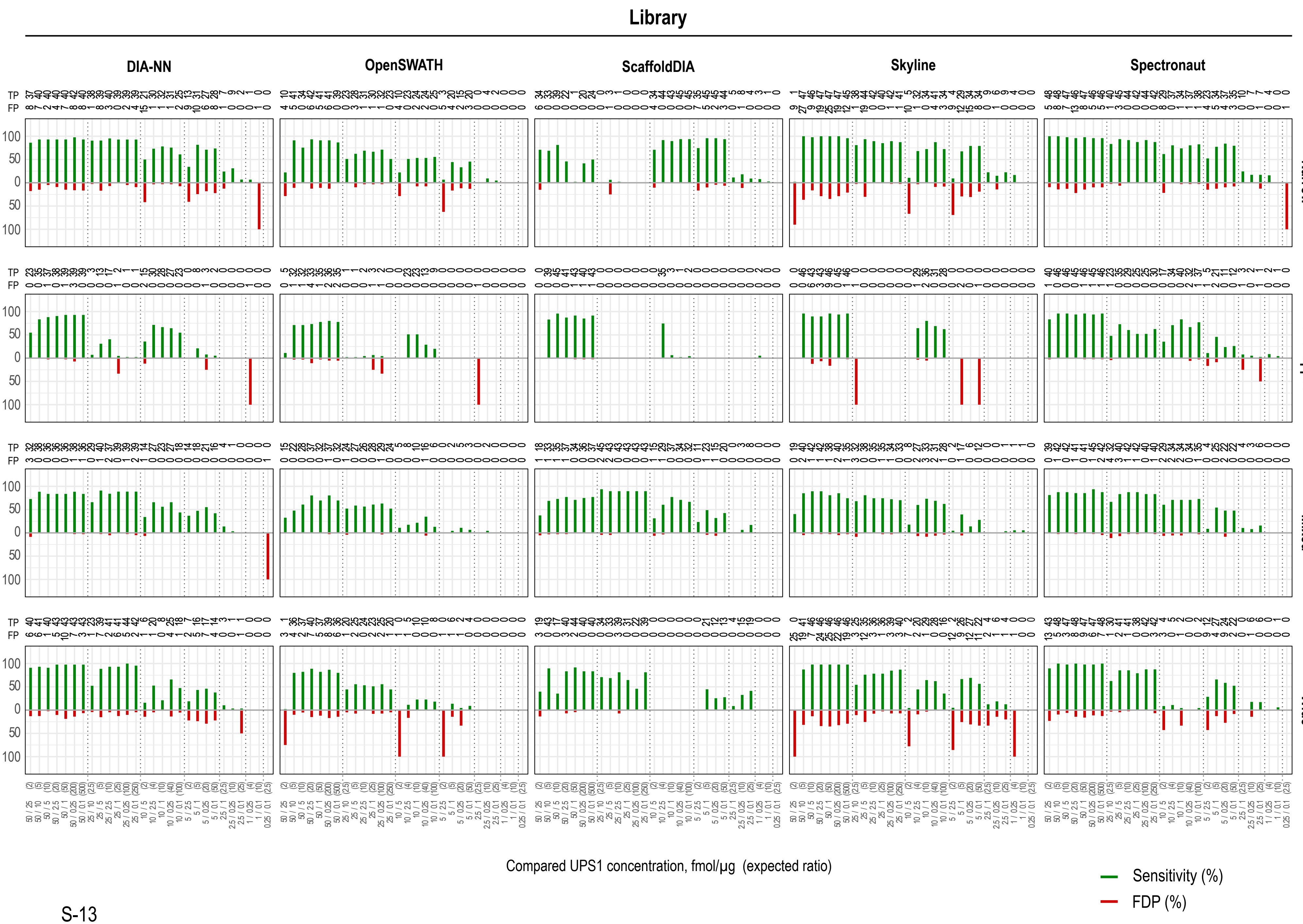
Library
DIA-NN
OpenSWATH
ScaffoldDIA
Skyline
Spectronaut
TP
FP
Narrow
Overlapped
Mixed
Wide
Compared UPS1 concentration, fmol/µg (expected ratio)
Sensitivity (%)
FDP (%)

**Differential analysis** - The sensitivity (*green*) and the false discovery proportion (FDP) (*red*) are shown for each pairwise comparison. A protein is considered regulated (or differentially expressed) if $|z| > 1.96$ and *q-value* < 0.05. Therefore, True Positives (TP) = Regulated UPS1 proteins; False Positives (FP) = Regulated *E. coli* proteins; True Negative (TN) = Non-regulated *E. coli* proteins and False Negatives (FN) = Non-regulated UPS1 proteins. *Sensitivity (%) = 100 * TP / (TP+FN)* and *FDP (%) = 100 * FP / (FP+TP)*.

### Supplementary Figure S7

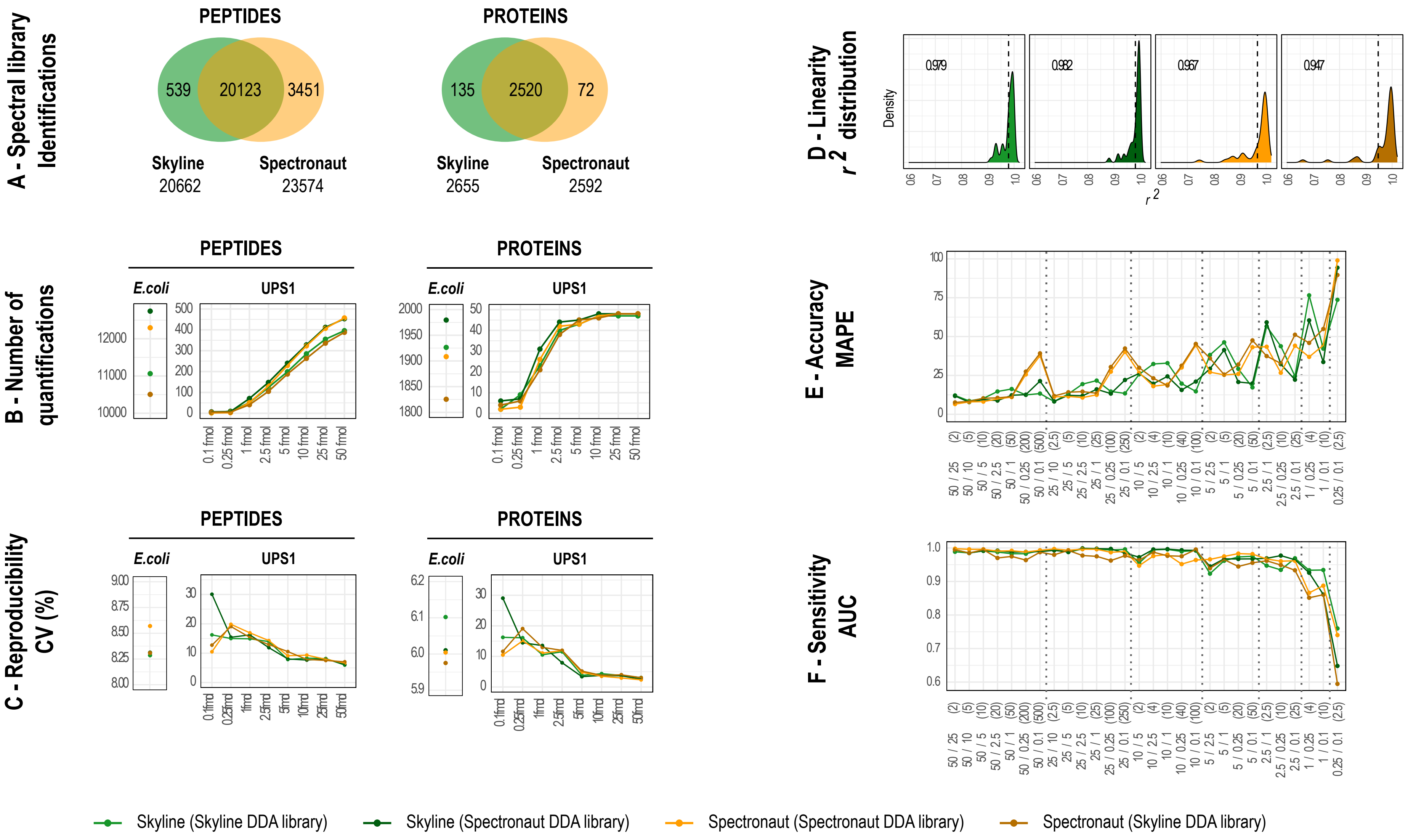


**Comparison of quantification performed by Skyline (*green*) and Spectronaut (*yellow)* using two different spectral libraries (*Library* mode), generated by Skyline or by Spectronaut, for *Narrow* acquisition scheme.** (**A**) Number of peptides and proteins contained in each spectral library; (**B**) Number of quantified peptides and proteins; (**C**) Coefficients of variation (%); (**D**) UPS1 coefficient of determination ($r^2$); (**E**) Mean absolute percentage error (MAPE) on Fold Change in pairwise comparisons and (**F**) Area under the curve (AUC) from ROC curves in pairwise comparisons.

### Supplementary Figure S8

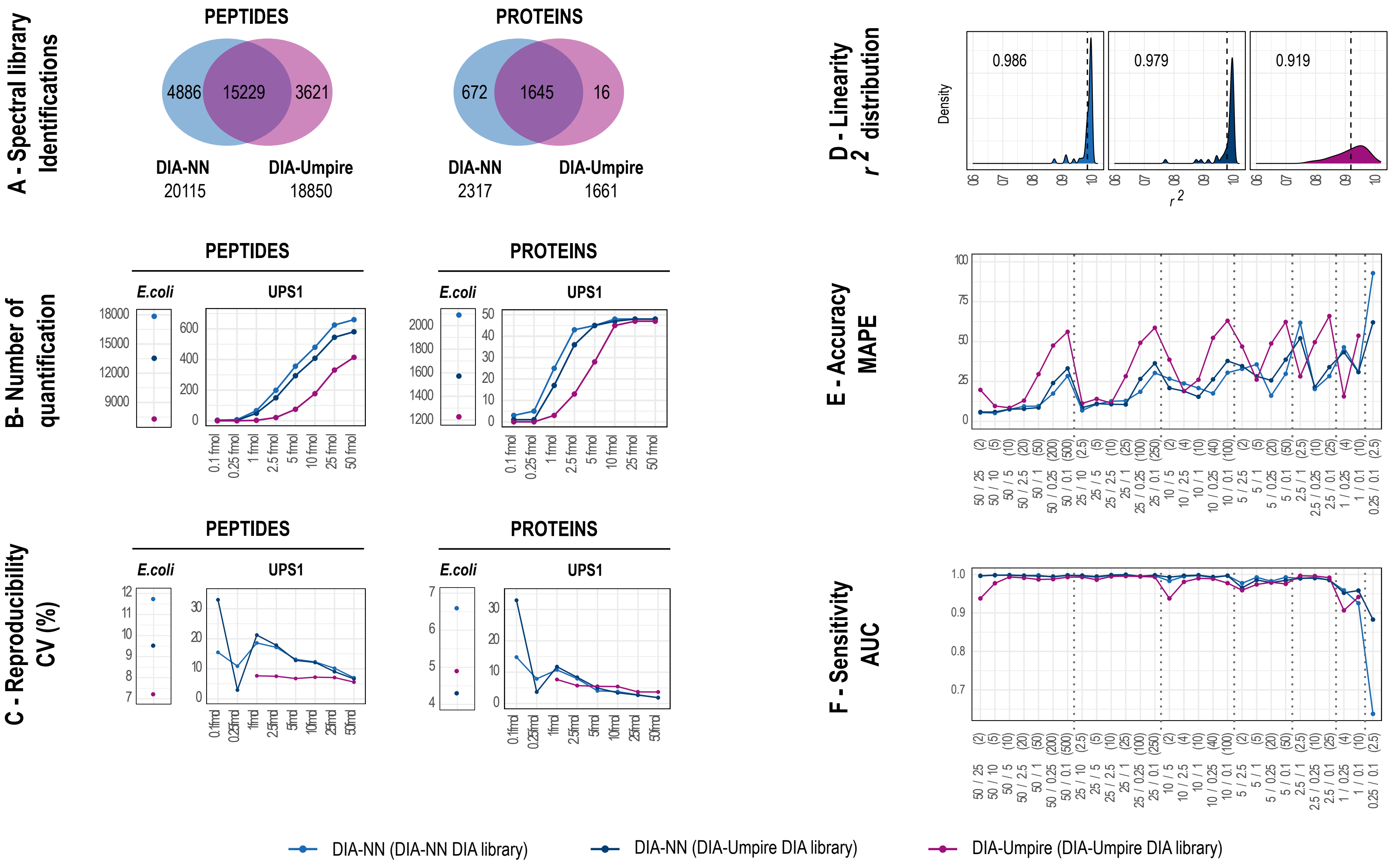


**Comparison of quantification performed with DIA-NN (*blue*) and DIA-Umpire (*purple*) with two different DIA spectral libraries (*FASTA* mode), generated by DIA-NN or DIA-Umpire (SE module + MSFragger), for *Narrow* acquisition scheme.** (**A**) Number of peptides and proteins contained in each spectral library; (**B**) Number of quantified peptides and proteins; (**C**) Coefficients of variation (%); (**D**) UPS1 coefficient of determination ($r^2$); (**E**) Mean absolute percentage error (MAPE) on Fold Change in pairwise comparisons and (**F**) Area under the curve (AUC) from ROC curves in pairwise comparisons.
